## Supplementary Information for "AIVE: accurate predictions of SARS-CoV-2 infectivity from comprehensive analysis"

Dongwan Hong, Ph.D.

Professor

College of Medicine, The Catholic University of Korea

222 Banpo-daero Seocho-gu, Seoul 06591, Republic of Korea

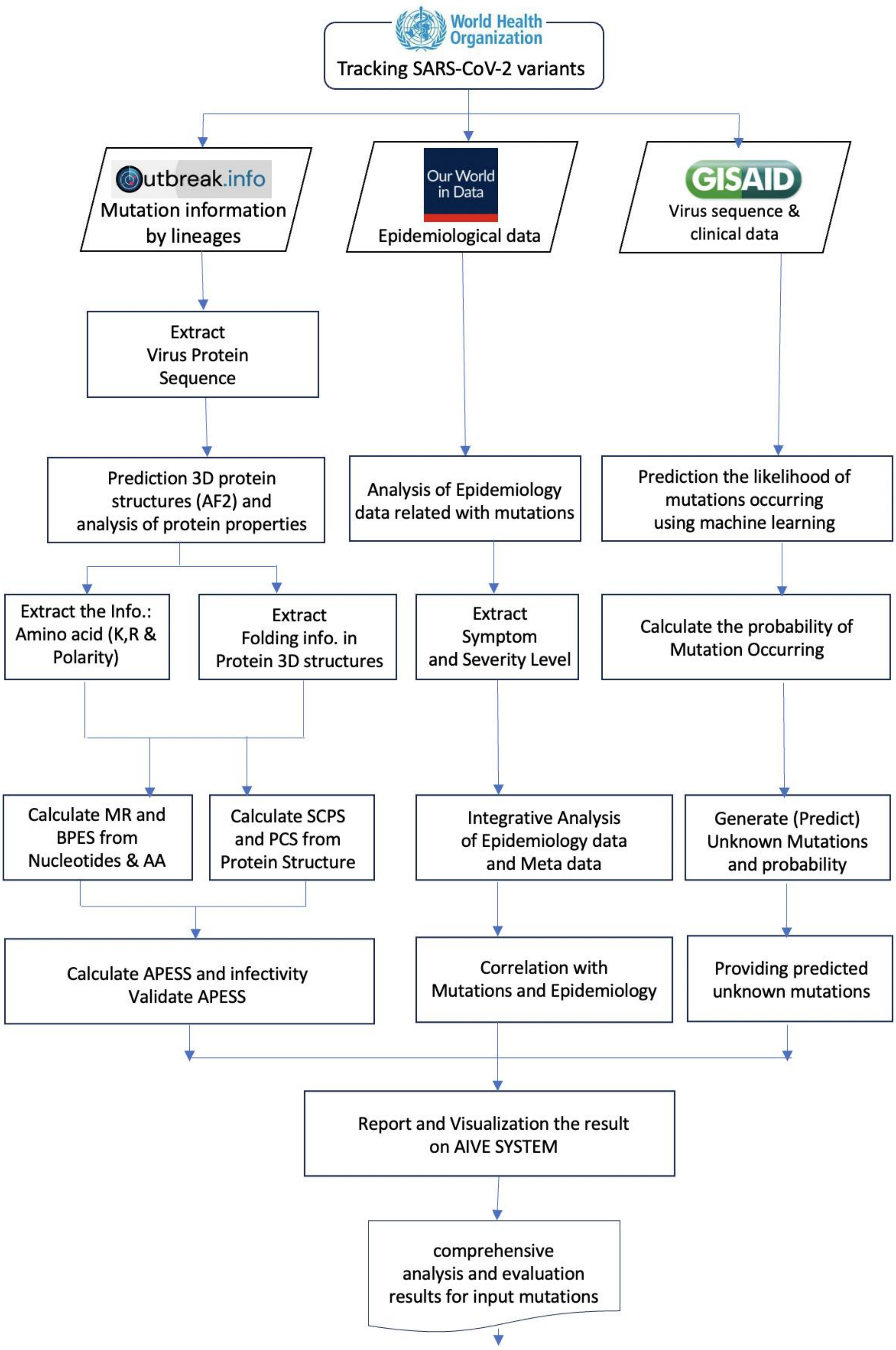

**Supplementary Figure 1. Research Overview**. AIVE can perform comprehensive analyses of SARS-CoV-2 mutations entered by User. From the World Health Organization (WHO), we obtained data from VOCs and VUMs and lineage mutation data from outbreak.info. Epidemiological data was obtained from OWID to provide the number of infections, deaths, vaccinations, and reproduction rate. From GISAID, virus sequence data and epidemiological data was collected. Through comprehensive analysis of databases, mutation prediction, evaluation, and epidemiological analysis based on mutation properties was carried out, and a web-based platform (AIVE) was created.

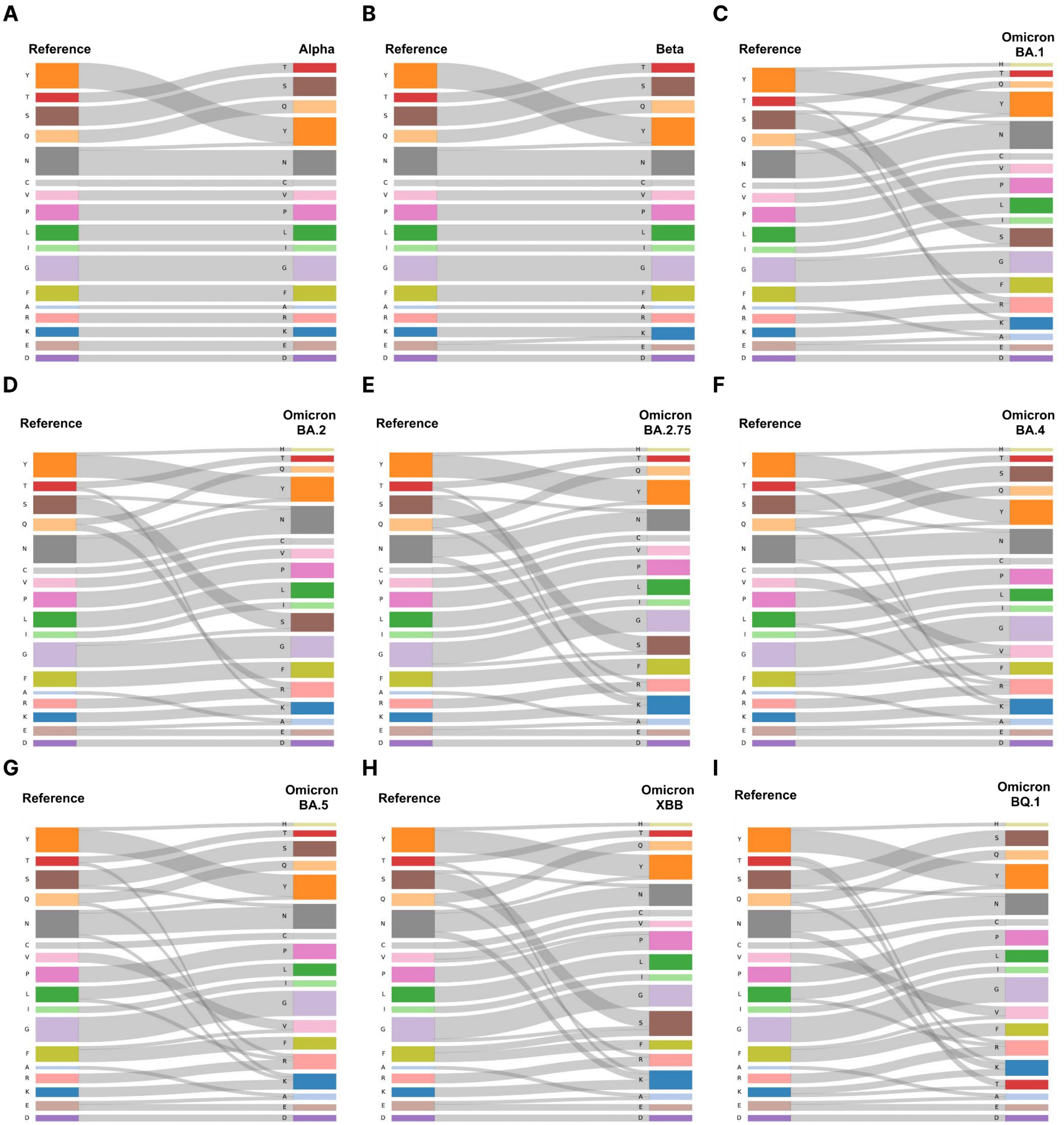

**Supplementary Figure 2. Amino acid substitutions in spike proteins from VOCs.** To show the changes of mutations according to evolution from Alpha to Omicron, we provide the Sankey diagram of each lineage. In VOCs (Alpha, Beta, Omicron BA.1, BA.2, BA.2.75, BA.4, BA.5, XBB, BQ.1), differences in amino acid substitutions were observed for each lineage compared to the reference. For Omicron sublineages, tyrosine (Y) is substituted for histidine (H). Also, there is an increase in lysine (K) and arginine (R) compared to the reference. The title of each lineage is as follows: **A)** Alpha, **B)** Beta, **C)** Omicron BA.1, **D)** Omicron BA.2, **E)** Omicron BA.2.75, **F)** Omicron BA.4, **G)** Omicron BA.5, **H)** Omicron XBB, and **I)** Omicron BQ.1

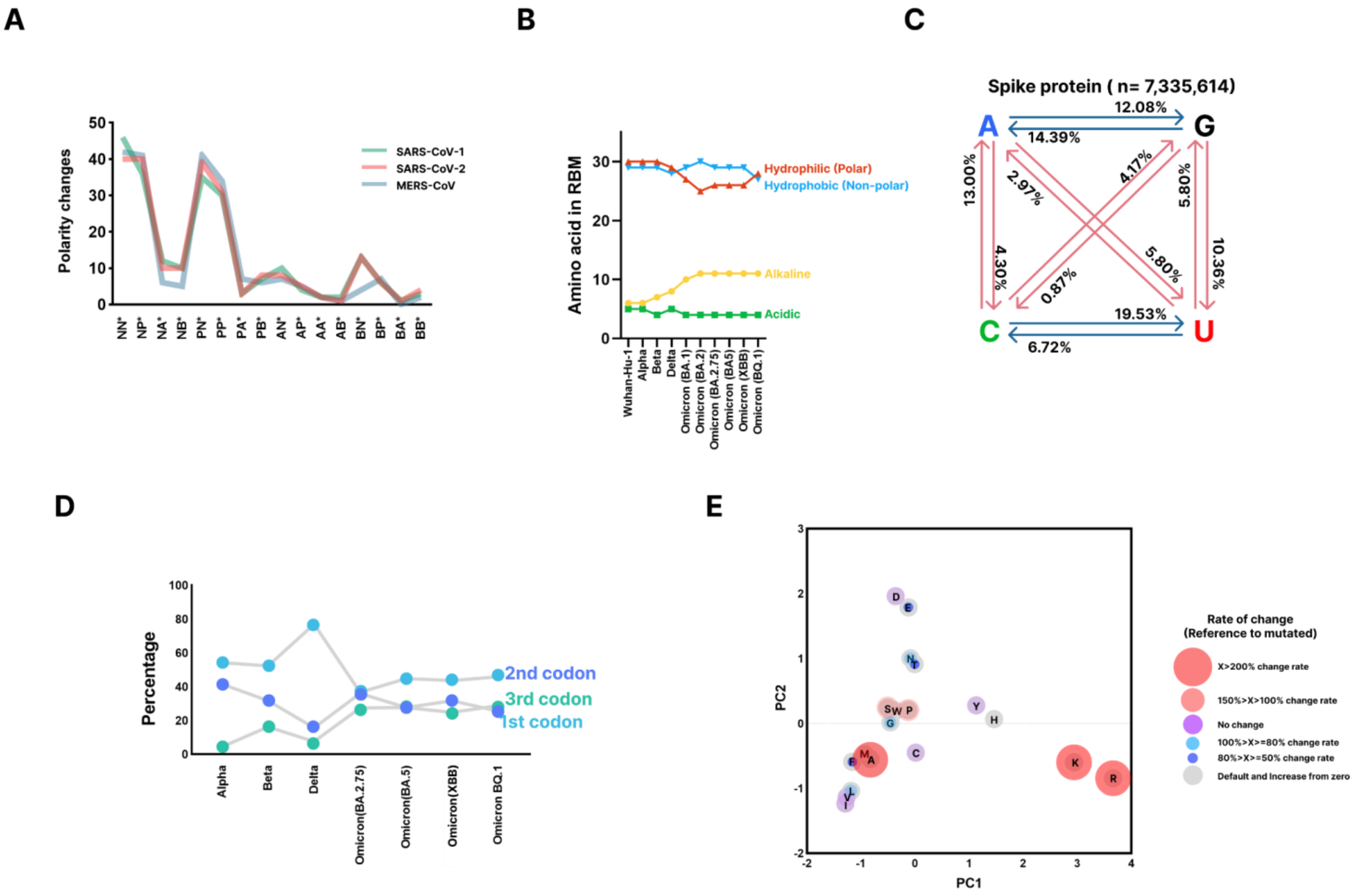

**Supplementary Figure 3. Polarity changes in the spike protein region of coronavirus and characteristics of RNA and amino acid levels of SARS-CoV-2. A)** The polarity of coronavirus: SARS-CoV-1, MERS-CoV, and SARS-CoV-2 are displayed. hydrophilic (N), hydrophobic (P), acidic (A), and basic (B). **B)** The number of amino acids in the RBM region for SARS-CoV-2 lineages (Wuhan-Hu-1, Alpha, Beta, Delta, BA.1, BA.2, BA.2.75, BA.4, XBB, and BQ.1) are displayed. As SARS-CoV-2 lineages progress from Wuhan-Hu-1 to Omicron, hydrophilic (polar) and hydrophobic (non-polar) amino acids are maintained at higher numbers. Meanwhile, there has been a slight increase in basic amino acids. There has not been a significant change in acidic amino acids. **C)** Transition and transversion rates for Adenine (A), Guanine (G), Cytosine (C), and Uracil (U) were counted and displayed (Red: transversions and Blue: transitions) for the spike protein of the 7,335,614 samples. **D)** We summarized the occurrences of SARS-CoV-2 mutations for the three codon positions comprising amino acids and investigated the mutation rate in the lineages (Alpha, Beta, Delta, Omicron (BA.2.75), Omicron (BA.4/BA.5), Omicron (XBB), Omicron BQ.1). The mutation rate for each codon position showed different values for each lineage. The mutation occurrence rate for the 2nd codon position was over 50% on average for all lineages. In the case of the Delta variant, the mutation rate for the 2nd codon position was the highest among all variants at 76.94%. **E)** The amino acid substitutions are displayed, and the size of the circle signifies the rate of change for the mutations against the reference. Lysine (K), arginine (R), and alanine (A) have the highest rate of change at over 200%.

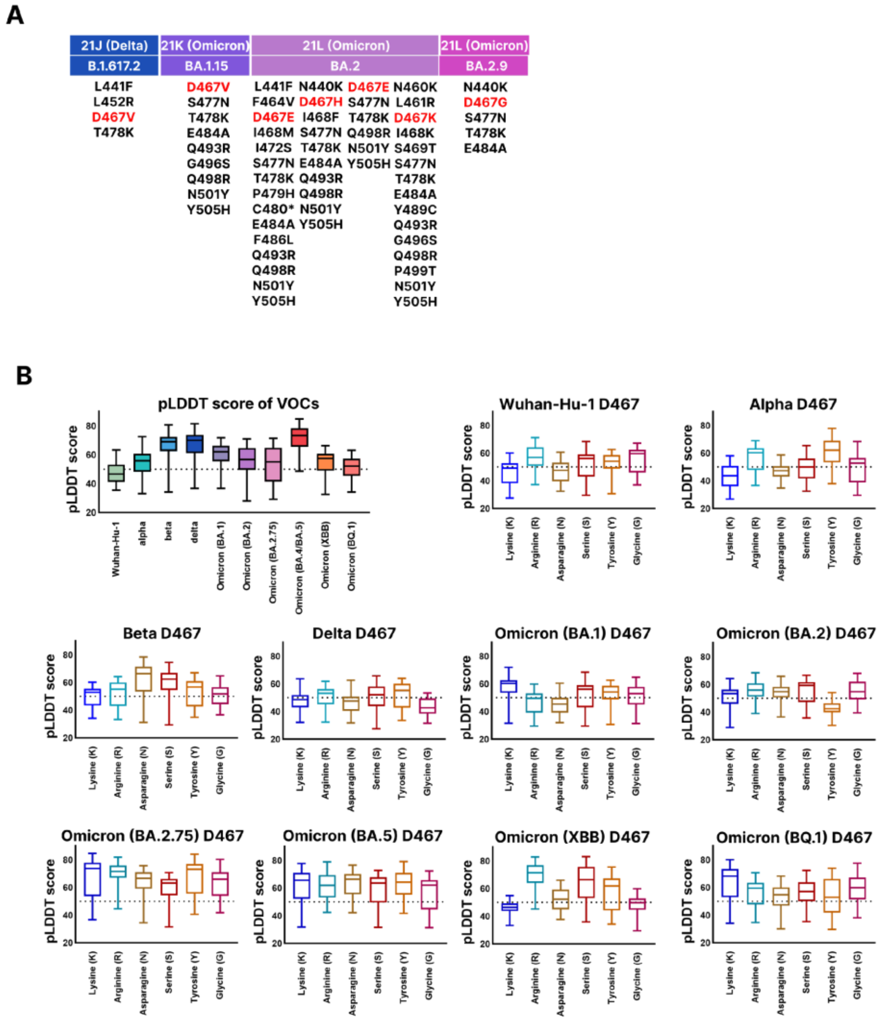

**Supplementary Figure 4. Presence of D467 mutation from GISAID and evaluation of protein structure due to D467. A)** Of the investigated sequences, there are very few instances of D467 amino acid substitutions. In the Delta variant, D467V was observed. In the Omicron BA.1.15 variant, D467V was observed. In the Omicron BA.2 variant, D467E, D467H, D467E, and D467K were observed. In the Omicron BA.2.9 variant, D467G was observed. **B**) The pLDDT scores are displayed on the y-axis for the graphs. Protein structure prediction of mutagenesis sequences at D467 showed that there was a general decrease in pLDDT score compared to VOCs. For SARS-CoV-2 variants, mutagenesis at D467 to lysine (K), arginine (R), glycine (G), tyrosine (Y), asparagine (N), serine (S) was carried out for Wuhan-Hu-1, Alpha, Beta, Delta, and Omicron (BA.1, BA.2, BA.2.75, BA.4/BA.5, XBB, BQ.1). We compared the pLDDT scores of the D467 mutagenesis variants against VOCs. For Wuhan-Hu-1 (47.96902778), Alpha (53.94541667), Delta (67.10597222), BA.1 (60.06305556), BA.2 (56.23902778), there was a general decrease in pLDDT score compared to VOCs. For Beta (65.915), BA.5 (70.96763889), BQ.1 (51.05027778), they showed similar pLDDT scores compared to the VOCs. The BA.2.75 (55.45305556) variant showed an increase in pLDDT score. However, for the pLDDT scores of all SARS-CoV-2 variants, the fluctuation of the confidence score was extreme and the PAE score was low as well.

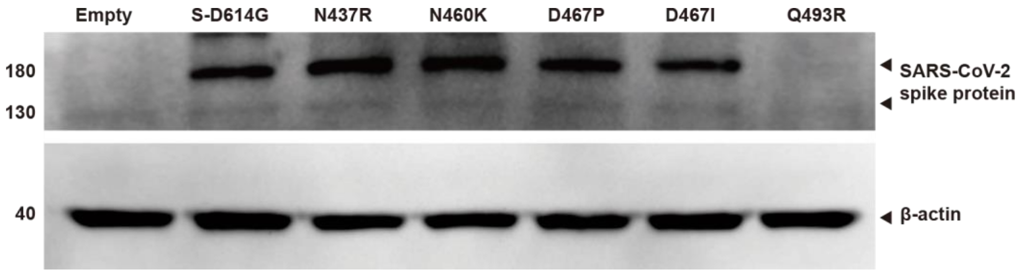

**Supplementary Figure 5. Identification of protein expression using western blotting**. We measured the expression of exogenous spike proteins through western blotting for SARS-CoV-2 spike protein mutations. To compare to the wild type (S-D614G), mutagenesis was done for N437R, N460K, D467P, D467I, and Q493R. Normalization was done with Beta-actin. The Beta actin bands are observed uniformly across the lanes for the vector, wild type (D614G), and mutagenesis sequences.

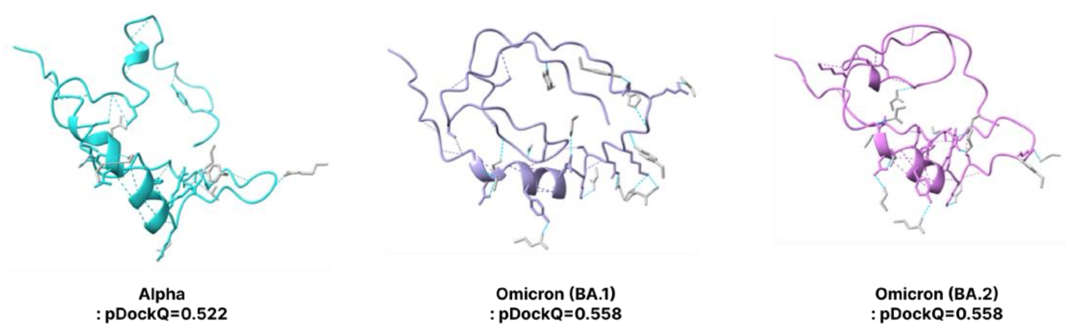

**Supplementary Figure 6. Prediction of spike protein 3D structures from Alpha, Omicron BA.1, Omicron BA.2.** The folding structures and pDockQ scores of Alpha, Omicron (BA.1), and Omicron (BA.2) are displayed. The pDockQ scores are 0.522, 0.558, and 0.558 respectively.

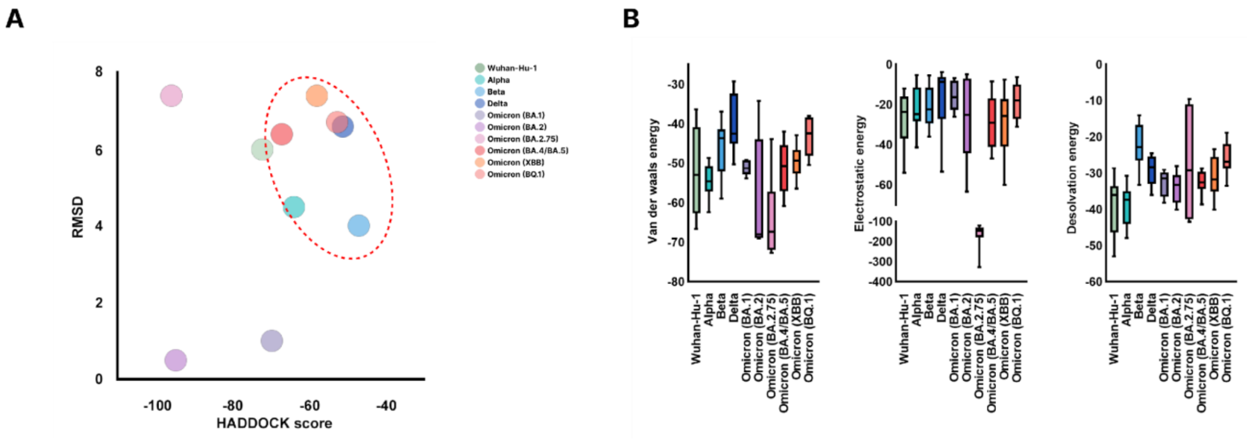

**Supplementary Figure 7. Binding affinity between SARS-CoV-2 and ACE2 receptor using HADDOCK. A**) The Root Mean Square Deviation (RMSD) and HADDOCK scores are displayed for SARS-CoV-2 lineages Wuhan-Hu-1, Alpha, Beta, Delta, Omicron (BA.1), Omicron (BA.2), Omicron (BA.2.75), Omicron (BA.4/BA.5), Omicron (XBB), and Omicron (BQ.1). The SARS-CoV-2 lineages are shown on the graph with corresponding colors. The SARS-CoV-2 lineages with both higher RMSD and HADDOCK scores are indicated with an encompassing red circle. **B**) For SARS-CoV-2 lineages Wuhan-Hu-1, Alpha, Beta, Delta, Omicron (BA.1), Omicron (BA.2), Omicron (BA.2.75), Omicron (BA.4/BA.5), Omicron (XBB), and Omicron (BQ.1), the van der waals energy, electrostatic energy, and desolvation energy are displayed in their respective graphs.

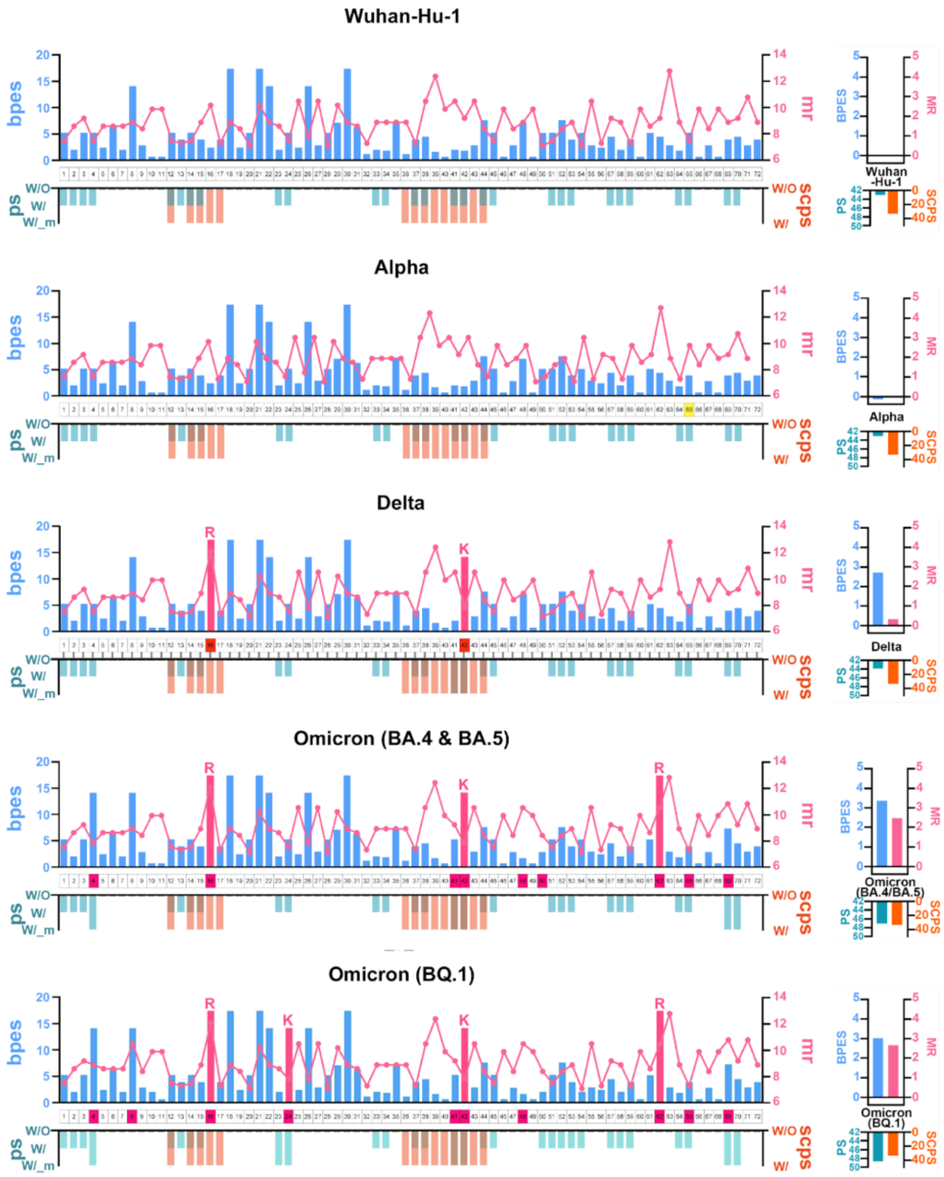

**Supplementary Figure 8. APESS evaluation of each section from VOCs.** For SARS-CoV-2 lineages Wuhan-Hu-1, Alpha, Delta, Omicron (BA.4/ BA.5), and Omicron (BQ.1), the SCPS, PCS, MR, and BPES scores are displayed. For SARS-CoV-2 lineages Wuhan-Hu-1, Alpha, Delta, Omicron (BA.4/ BA.5), and Omicron (BQ.1), the SCPS, PCS, MR, and BPES scores were calculated. For the RBM 72 positions, a comprehensive evaluation was carried out. The colors in the figure correspond with the following: sky blue: BPES, pink: MR, khaki: PCS, orange: SCPS, and pink box: lysine (K) and arginine (R). As the lineages progressed to Omicron, SCPS, PCS, MR and BPES scores were increased. Finally, APESS has higher score in the most infectious lineage.

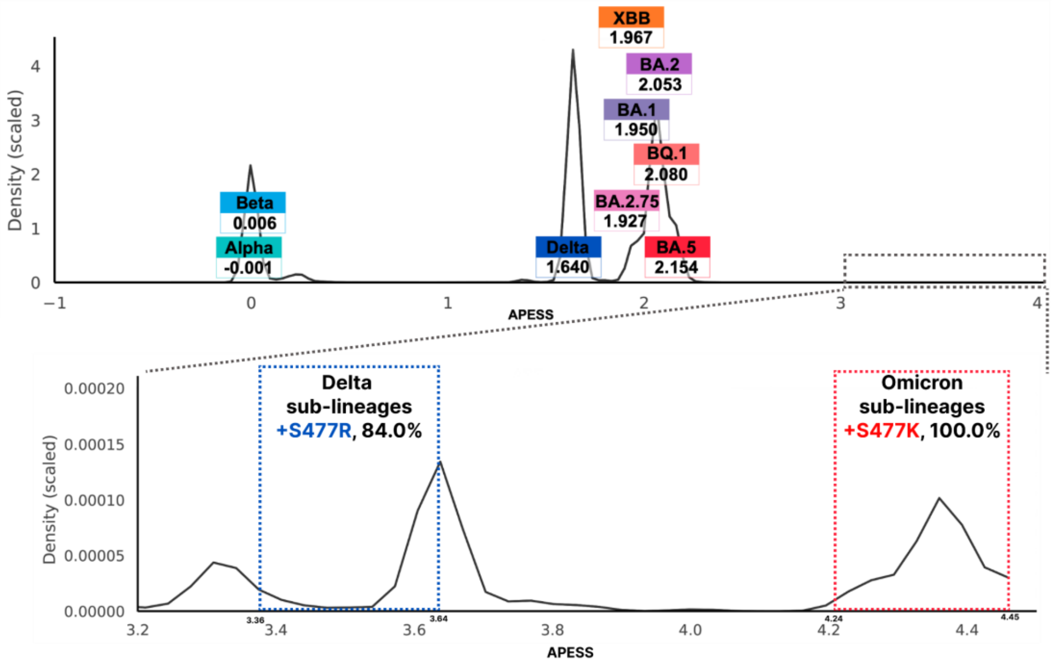

**Supplementary Figure 9. APESS distribution from viral sequences and APESS scores of VOCs from GISAID.** We measured the APESS score distribution for total 7,335,614 sublineages. The graph was identical to the one created from the randomly selected 30,000 sublineages. We verified four components centered at 0.0236, 1.6062, 1.6404, and 2.0627. Other than the four components, density was very low. High APESS scores were observed within the low-density region, specifically for S477K and S477R mutations. In the Delta variant, amino acid substitution from Serine (S) to Arginine (R) was observed with one nucleotide change at the final codon. In the Omicron variant, amino acid substitution from Serine (S) to Lysine (K) was observed with two nucleotide changes. In other SARS-CoV-2 variants, amino acid substitution from Serine (S) to Aspartic acid (N) with one nucleotide change at the middle codon.

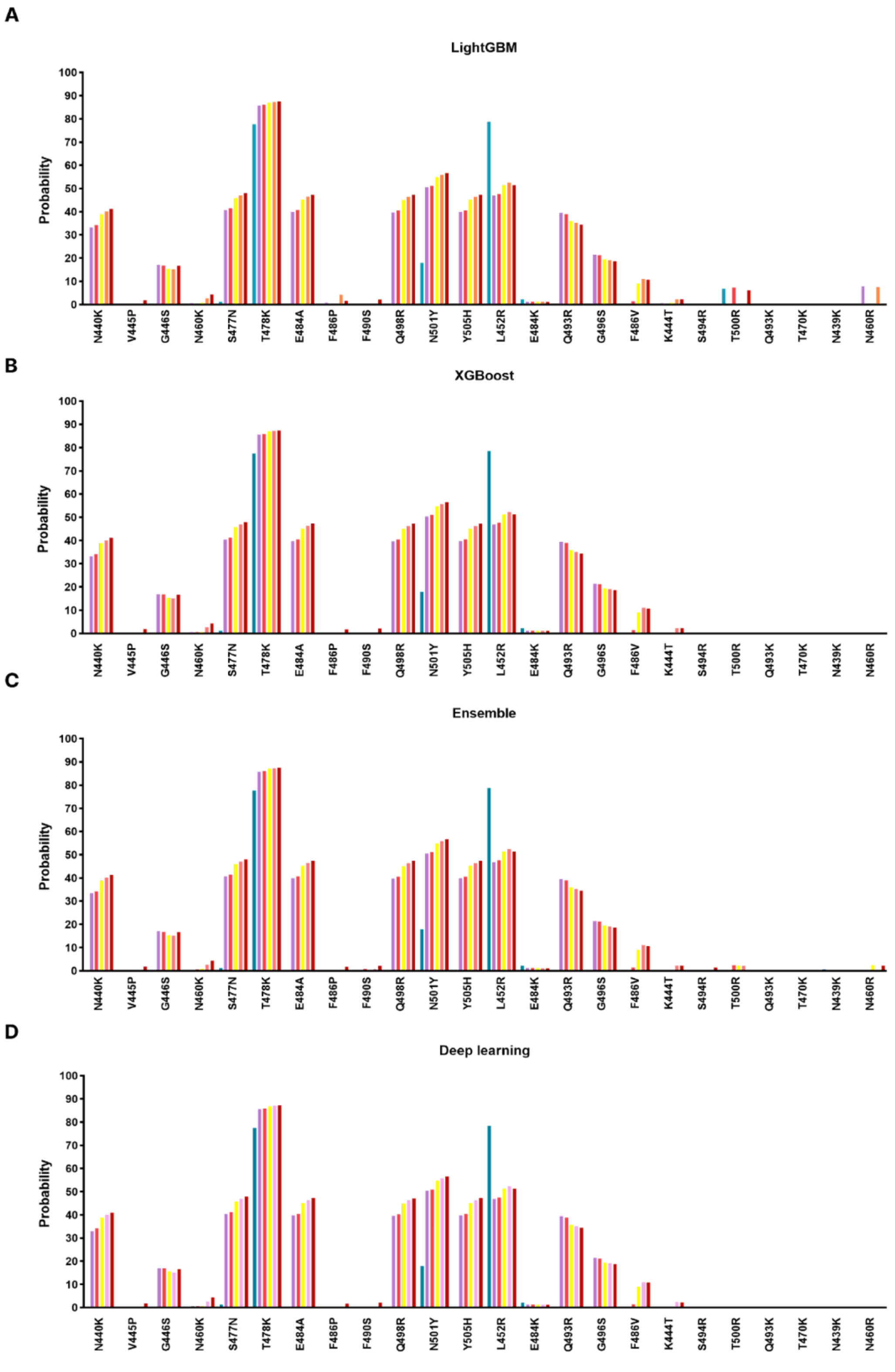

**Supplementary Figure 10. Prediction results of notable mutations in SARS-CoV-2 using ML/DL.** For mutations occurring in SARS-CoV-2 lineages (N440K, V445P, G446S, N460K, S477N, T478K, E484A, F486P, F490S, Q498R, N501Y, Y505H, L452R, E484K, Q493R, G496S, F486V, K444T) and mutations evaluated through APESS (N477R, N477K, N439R, Y501R, N437R, S438R, S459R, S469R, S494R, T470R, T500R, Q493K, T470K, T500K, S469K, S494K, Y501K, N437K, N439K, N460R, S438K, S459K, N460K, Q493R), artificial intelligence learning models were used to evaluate probability. LightGBM, XGBoost, Ensemble and deep learning methods showed similar results to Random Forest.

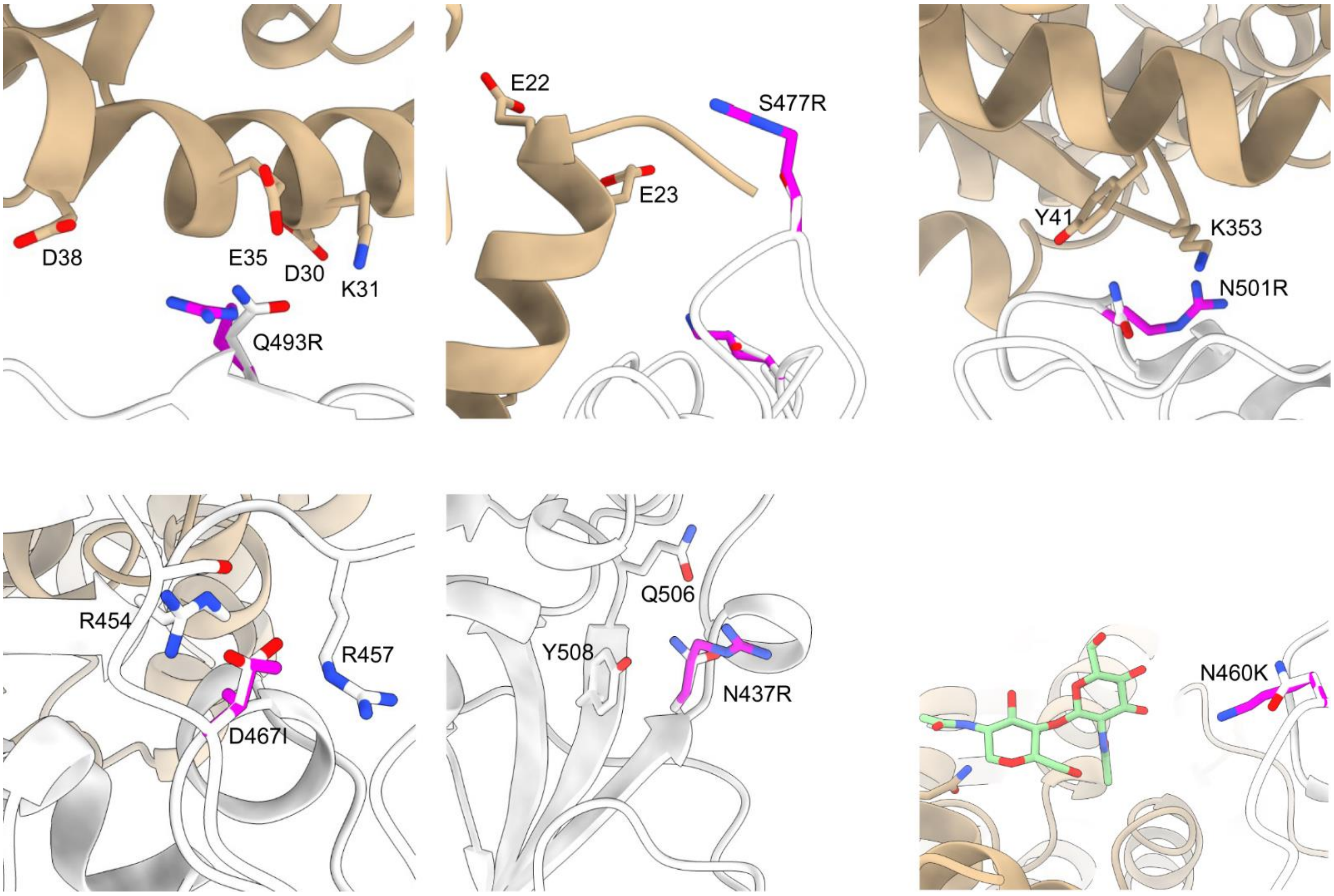

**Supplementary Figure 11. Protein structure evaluation of D467.** Close-up views of the mutated RBD residues and their adjacent residues. ACE2 and RBD are color-coded in wheat and silver cartoons, while the original and mutated RBD residues are represented as white and magenta sticks. Mutagenesis was carried out using PyMol, and all figure images were constructed using ChimeraX-1.5.

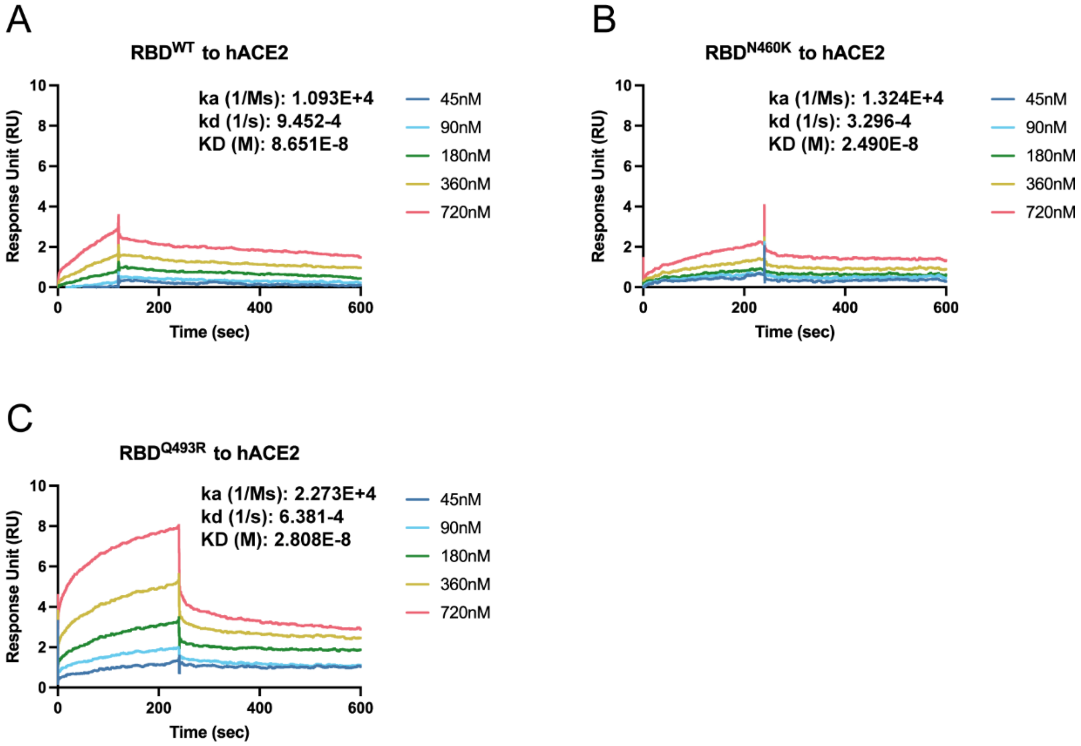

**Supplementary Figure 12. Binding affinity measurement of N460K and Q493R RBD variants.** surface plasmon resonance results showing binding affinity between RBD wild type (A), N460K (B), and Q493 (C) toward human ACE2. The hACE2 protein was used as ligand and all experiments were conducted on the same sensor chip.

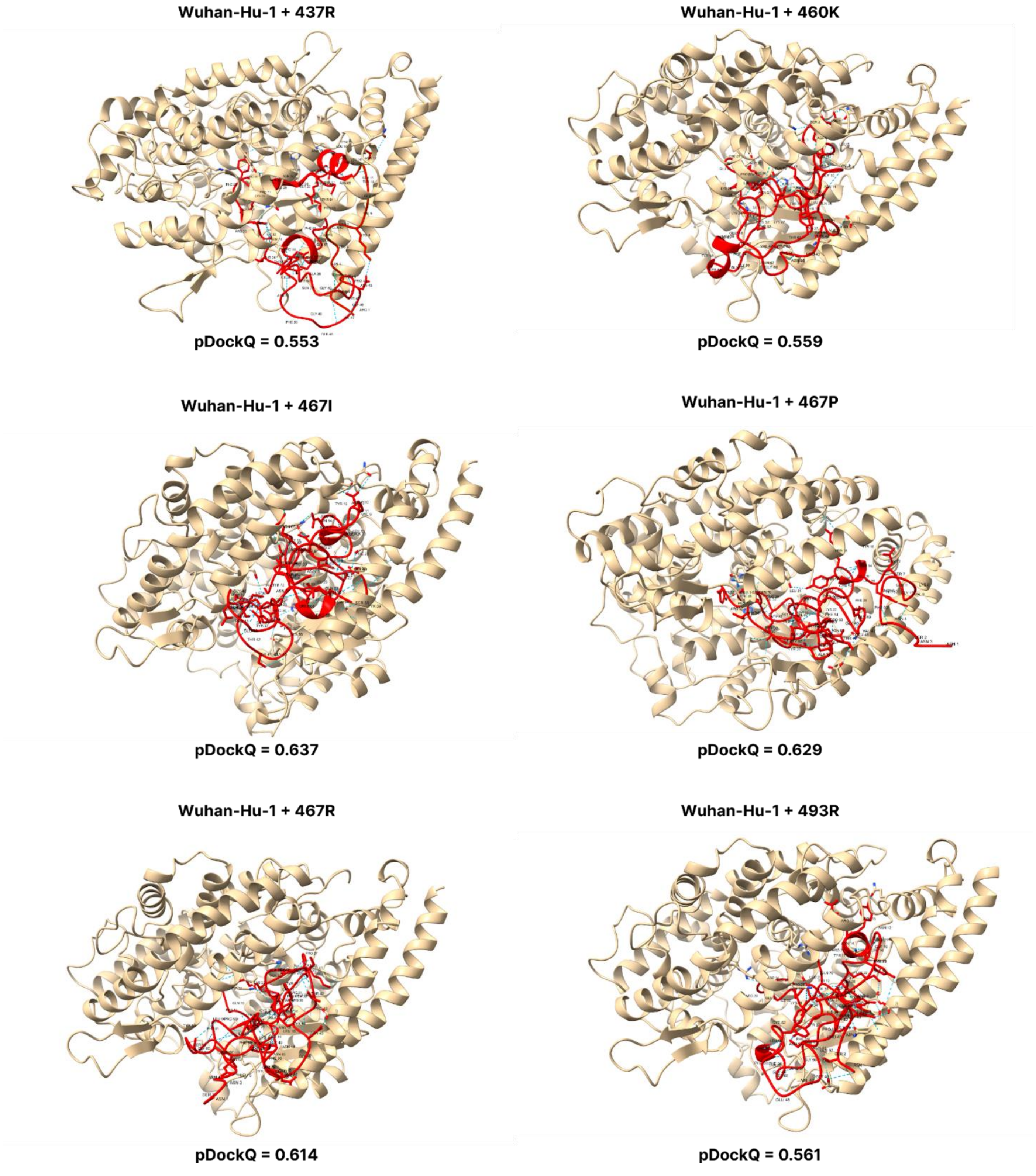

**Supplementary Figure 13. pDockQ scores of Wuhan-Hu-1 mutations.** pDockQ scores of lineages with the Wuhan-Hu-1 as the backbone, with each mutation N437R, N 460K, D467I, D467P, D467R, and Q493R added.

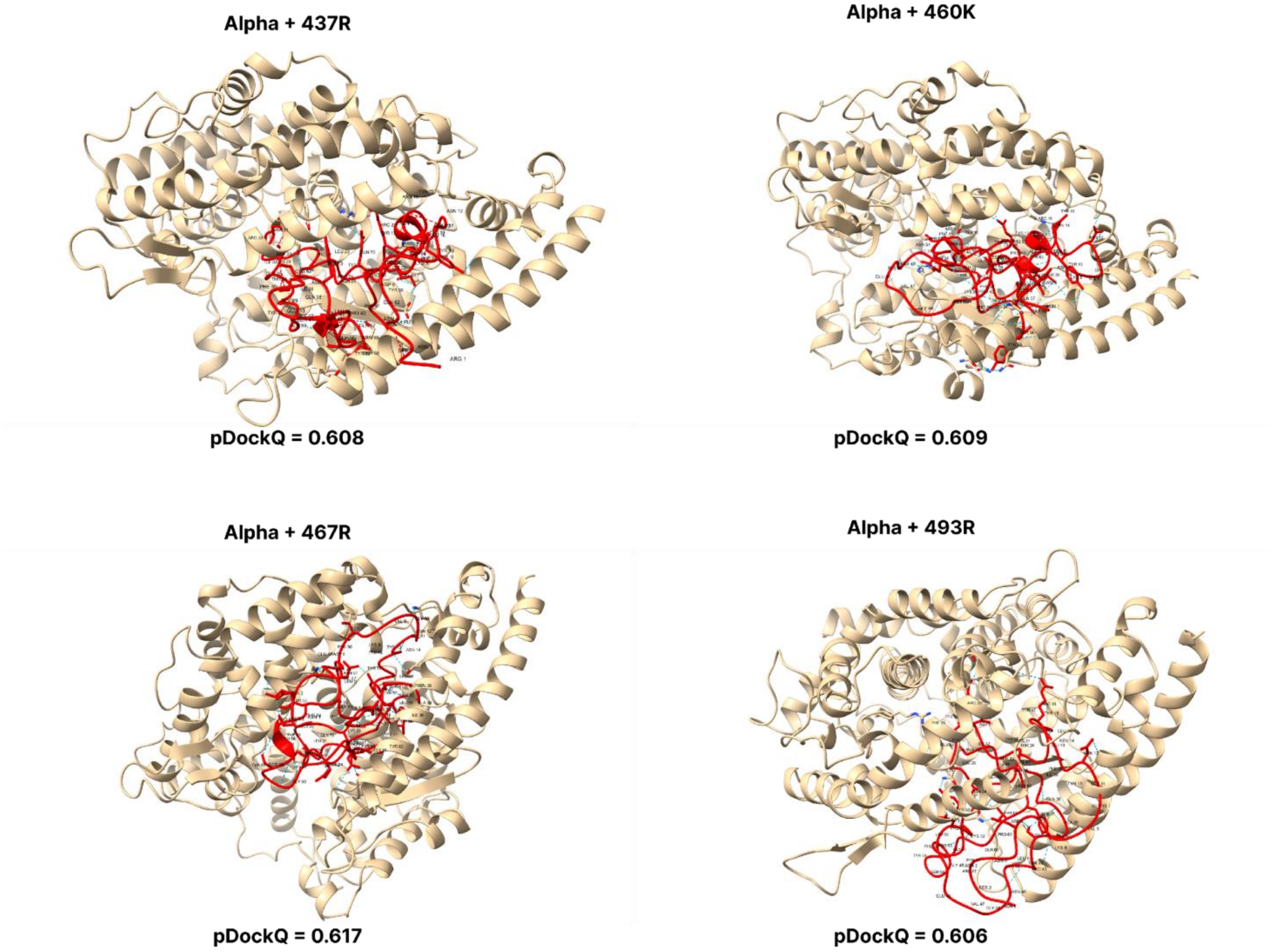

**Supplementary Figure 14. pDockQ scores of Alpha mutations.** pDockQ scores of lineages with the alpha lineages as the backbone, with each mutation N437R, N 460K, D467R, and Q493R added.

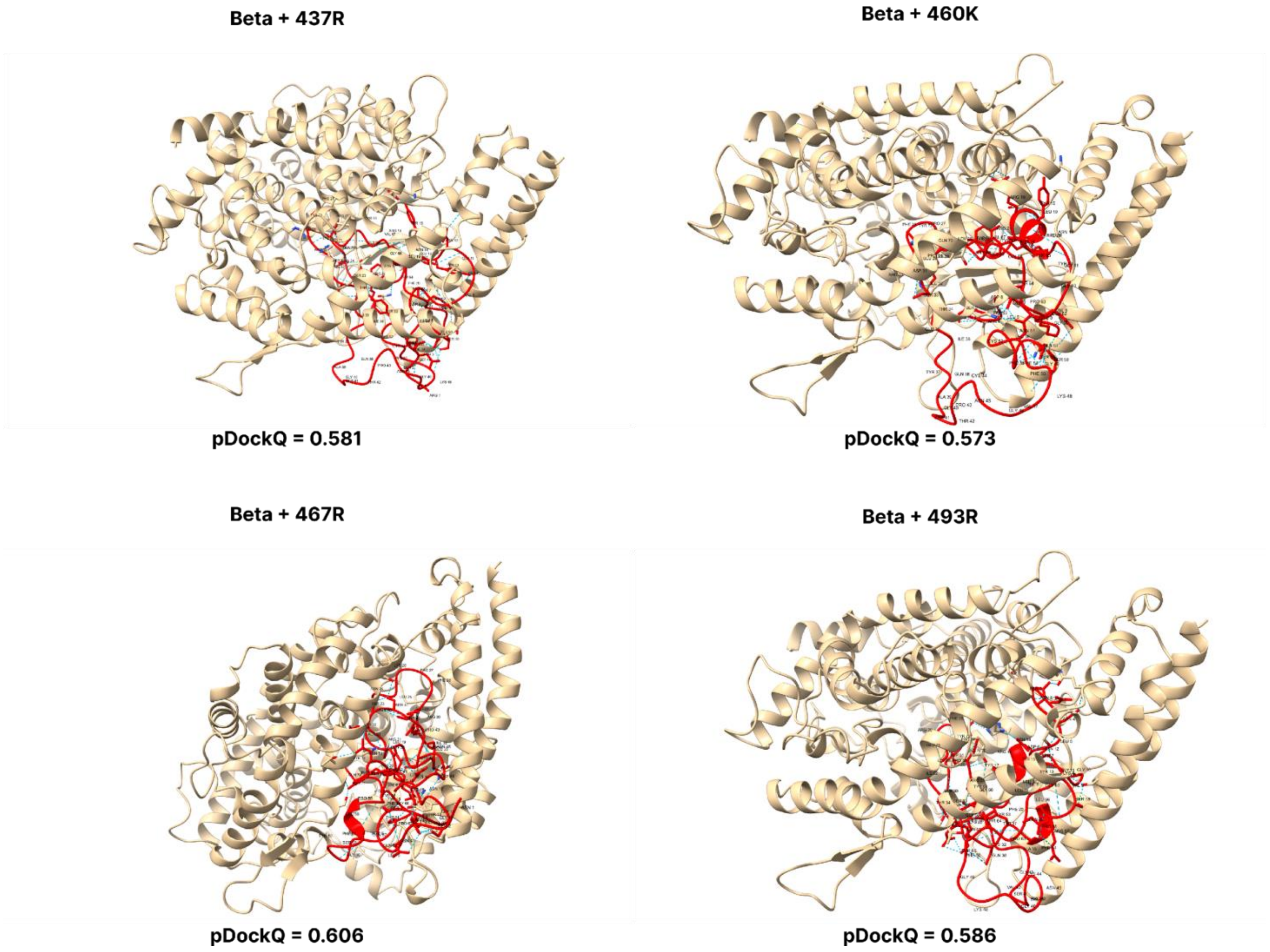

**Supplementary Figure 15. pDockQ scores of Beta mutations.** pDockQ scores of lineages with the beta lineages as the backbone, with each mutation N437R, N 460K, D467R, and Q493R added.

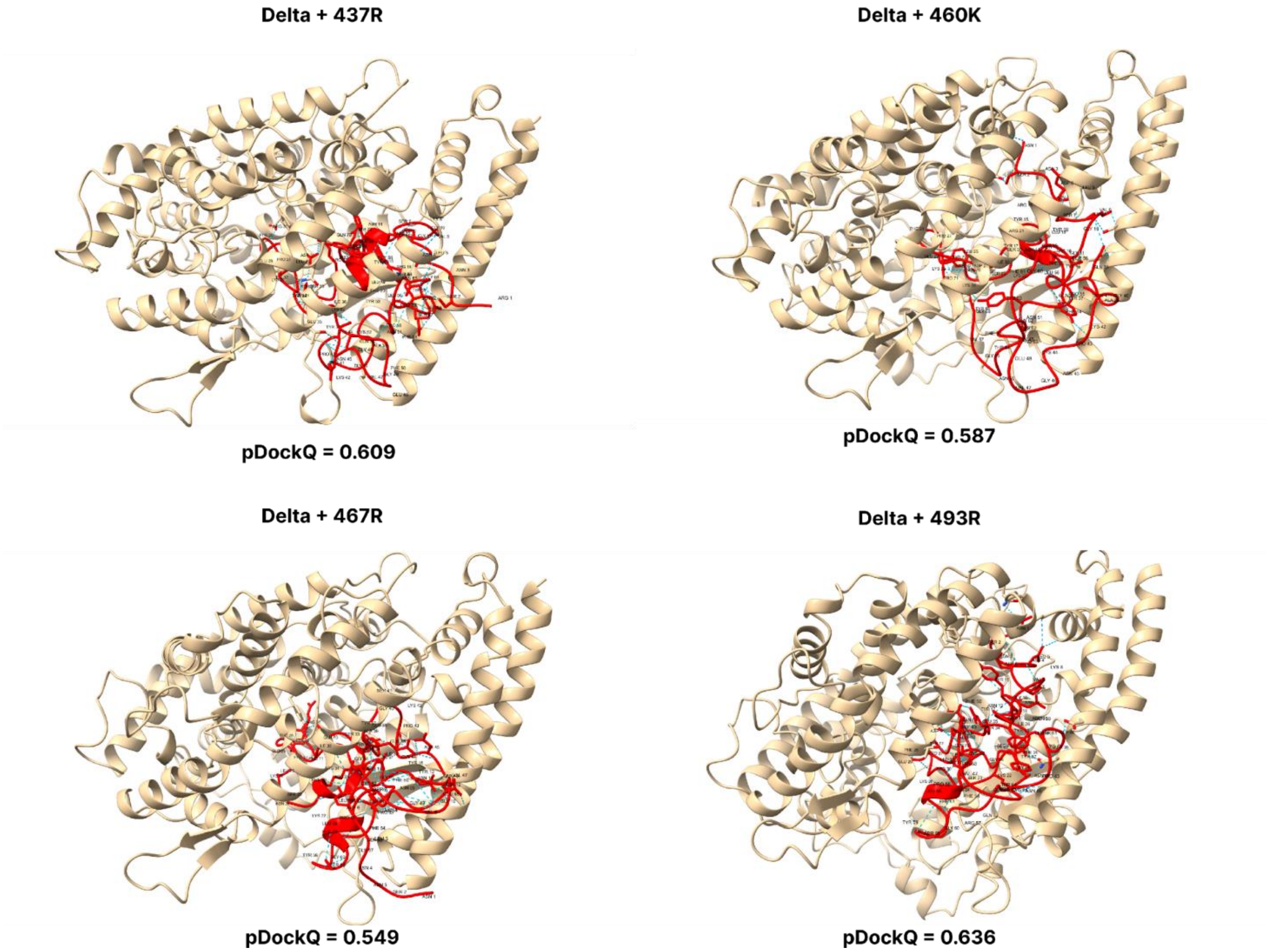

**Supplementary Figure 16. pDockQ scores of Delta mutations.** pDockQ scores of lineages with the delta lineages as the backbone, with each mutation N437R, N 460K, D467R, and Q493R added.

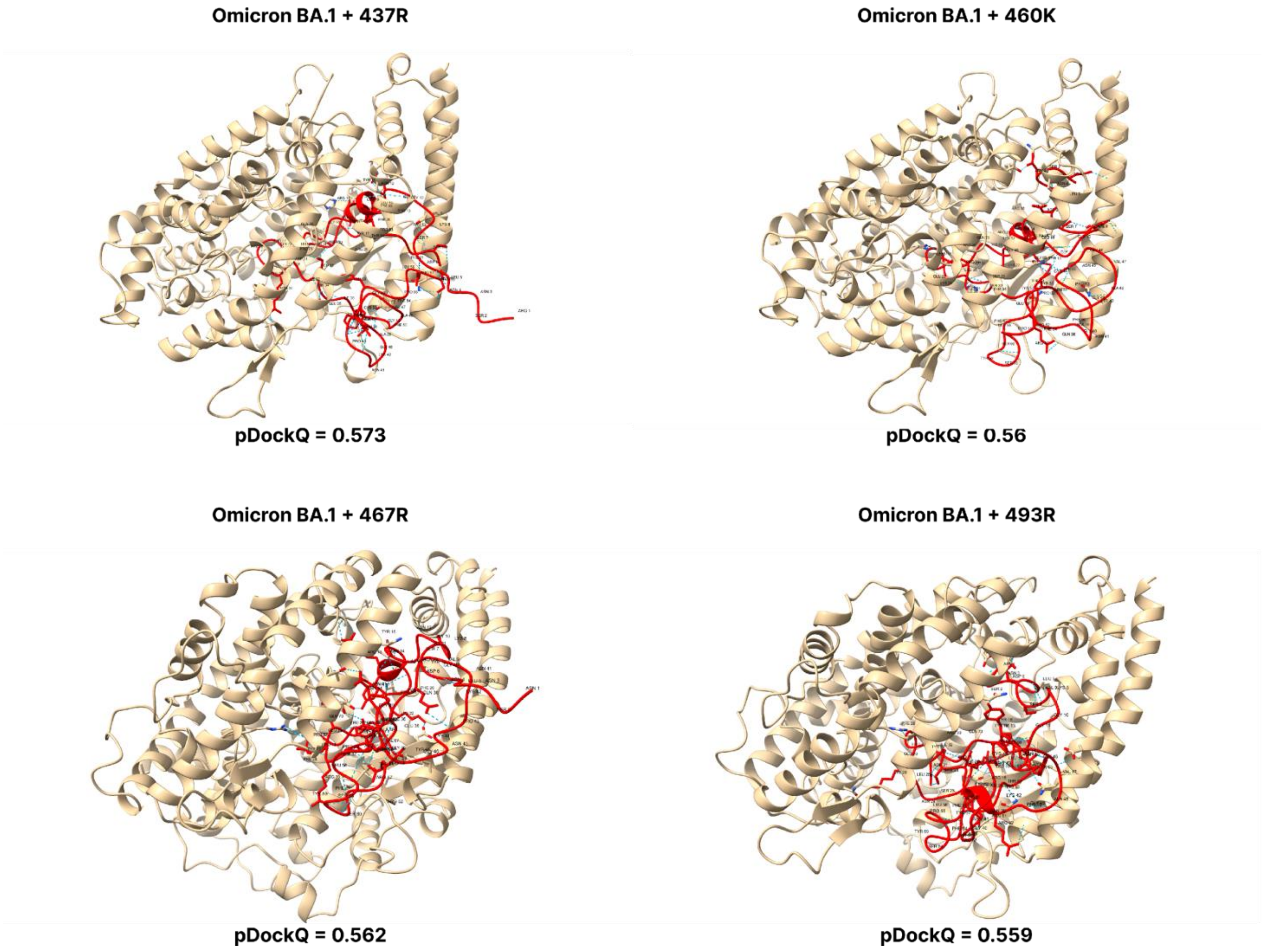

**Supplementary Figure 17. pDockQ scores of BA.1 mutations.** pDockQ scores of lineages with the omicron BA.1 as the backbone, with each mutation N437R, N 460K, D467R, and Q493R added.

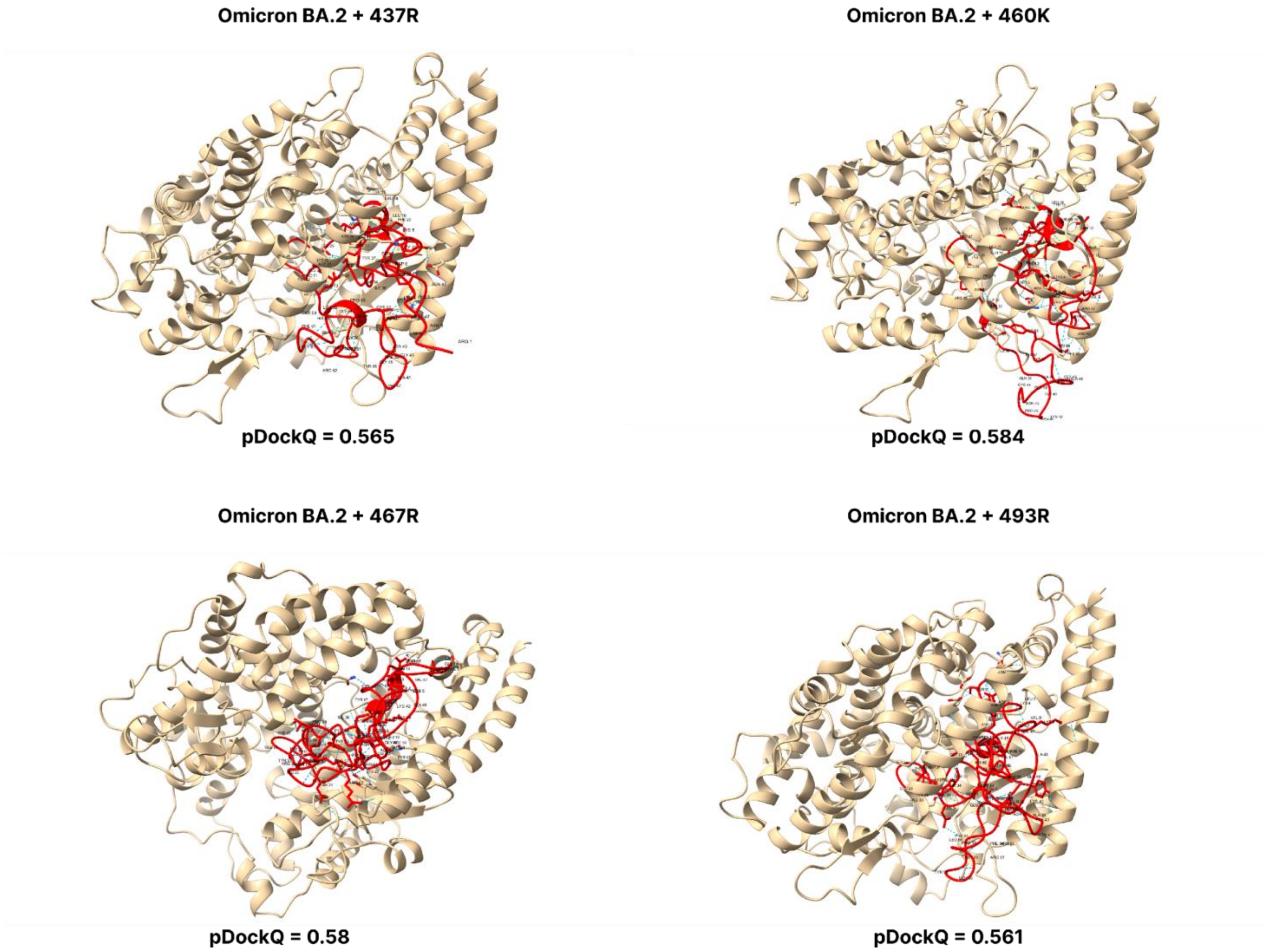

**Supplementary Figure 18. pDockQ scores of BA.2 mutations.** pDockQ scores of lineages with the omicron BA.2 lineages as the backbone, with each mutation N437R, N460K, D467R, and Q493R added.

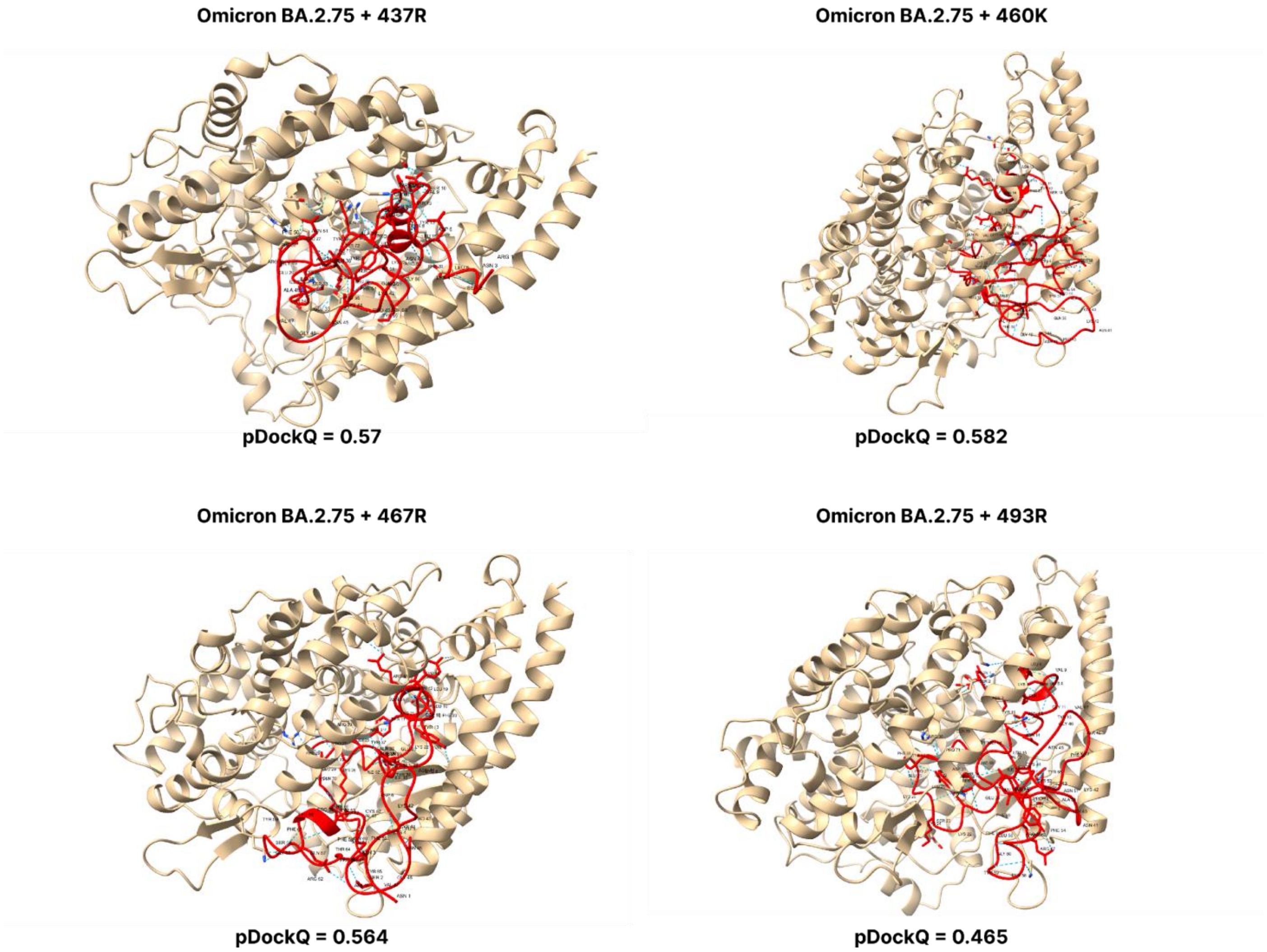

**Supplementary Figure 19. pDockQ scores of BA.2.75 mutations.** pDockQ scores of lineages with the omicron BA.2.75 lineages as the backbone, with each mutation N437R, N460K, D467R, and Q493R added.

**Omicron BA.4 + 437R**

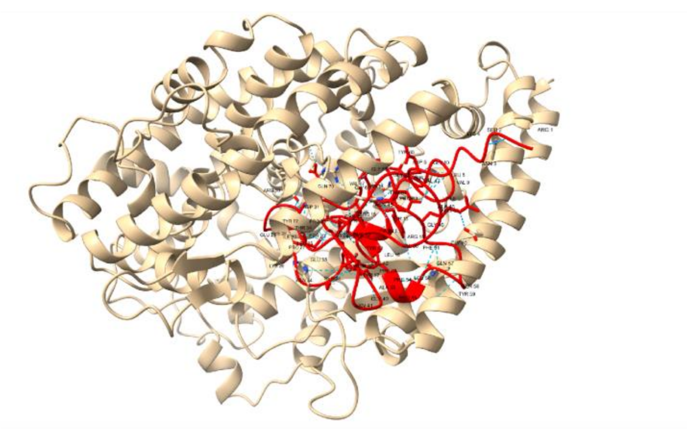

**pDockQ = 0.615**

**Omicron BA.4 + 460K**

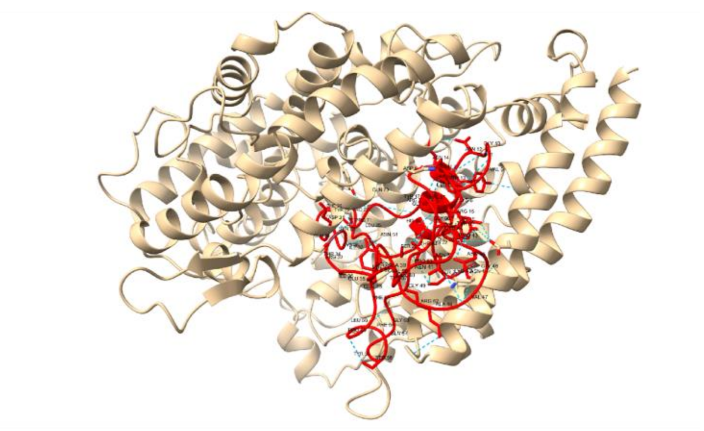

**pDockQ = 0.566**

**Omicron BA.4 + 467R**

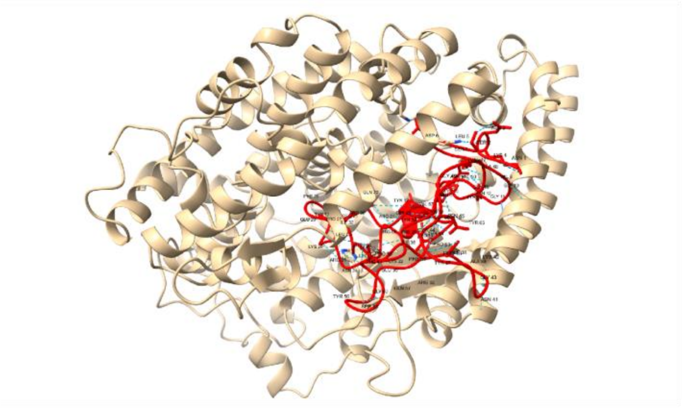

**pDockQ = 0.603**

**Omicron BA.4 + 493R**

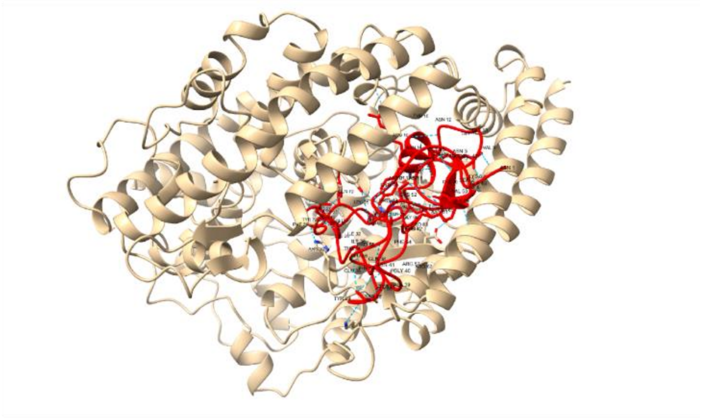

**pDockQ = 0.499**

**Supplementary Figure 20. pDockQ scores of BA.4 mutations.** pDockQ scores of lineages with the omicron BA.2.75 lineages as the backbone, with each mutation N437R, N460K, D467R, and Q493R added

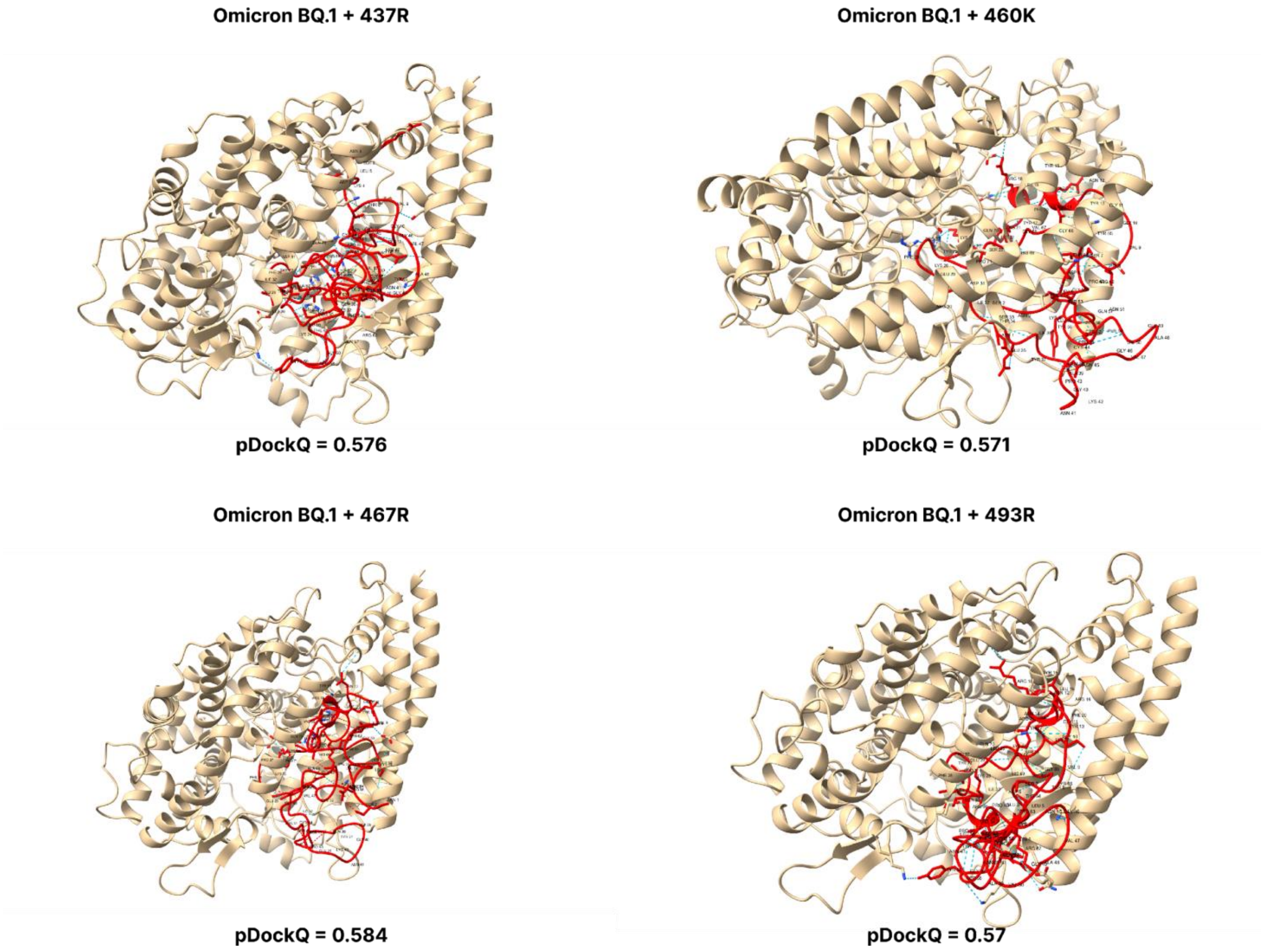

**Supplementary Figure 21. pDockQ scores of BQ.1 mutations.** pDockQ scores of lineages with the omicron BQ1 lineages as the backbone, with each mutation N437R, N460K, D467R, and Q493R added

**Omicron XBB + 437R**

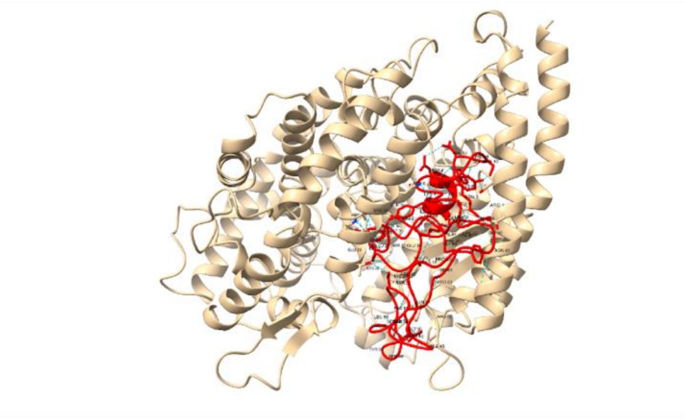

**pDockQ = 0.566**

**Omicron XBB + 460K**

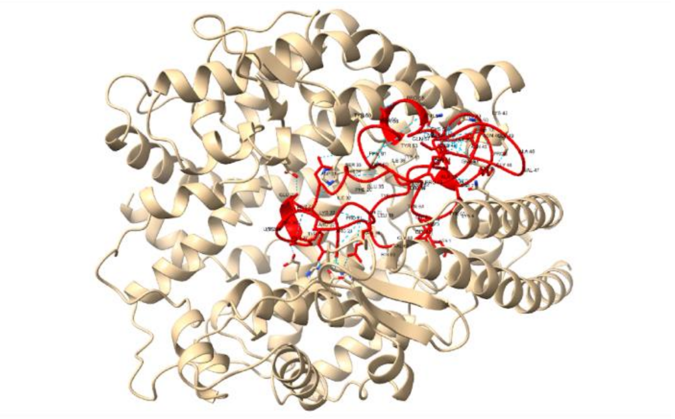

**pDockQ = 0.571**

**Omicron XBB + 467I**

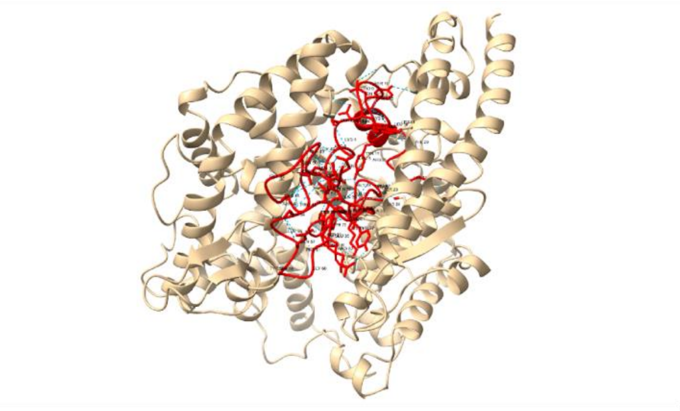

**pDockQ = 0.556**

**Omicron XBB + 467P**

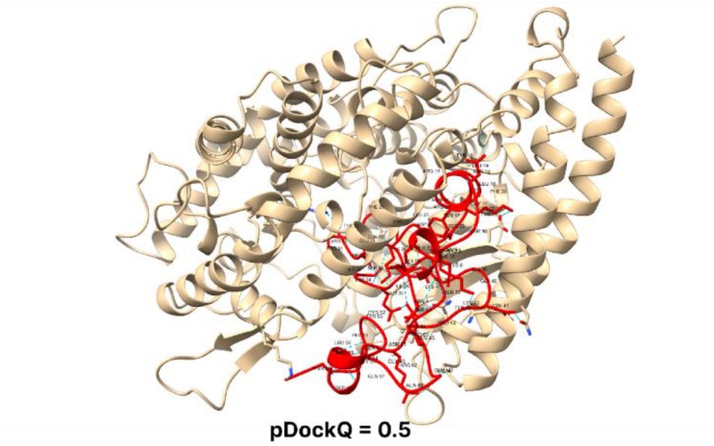

**pDockQ = 0.5**

**Omicron XBB + 467R**

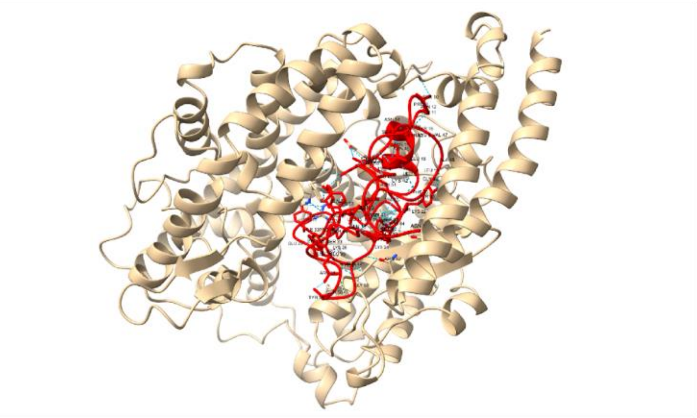

**pDockQ = 0.591**

**Omicron XBB + 493R**

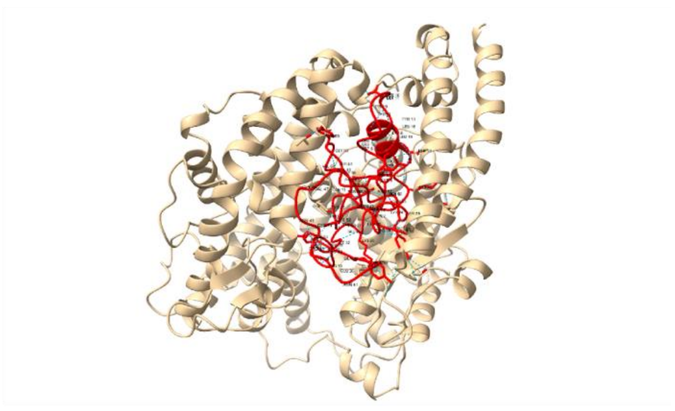

**pDockQ = 0.585**

**Supplementary Figure 22. pDockQ scores of XBB mutations.** pDockQ scores of lineages with the omicron BA.2.75 lineages as the backbone, with each mutation N437R, N460K, D467R, and Q493R added

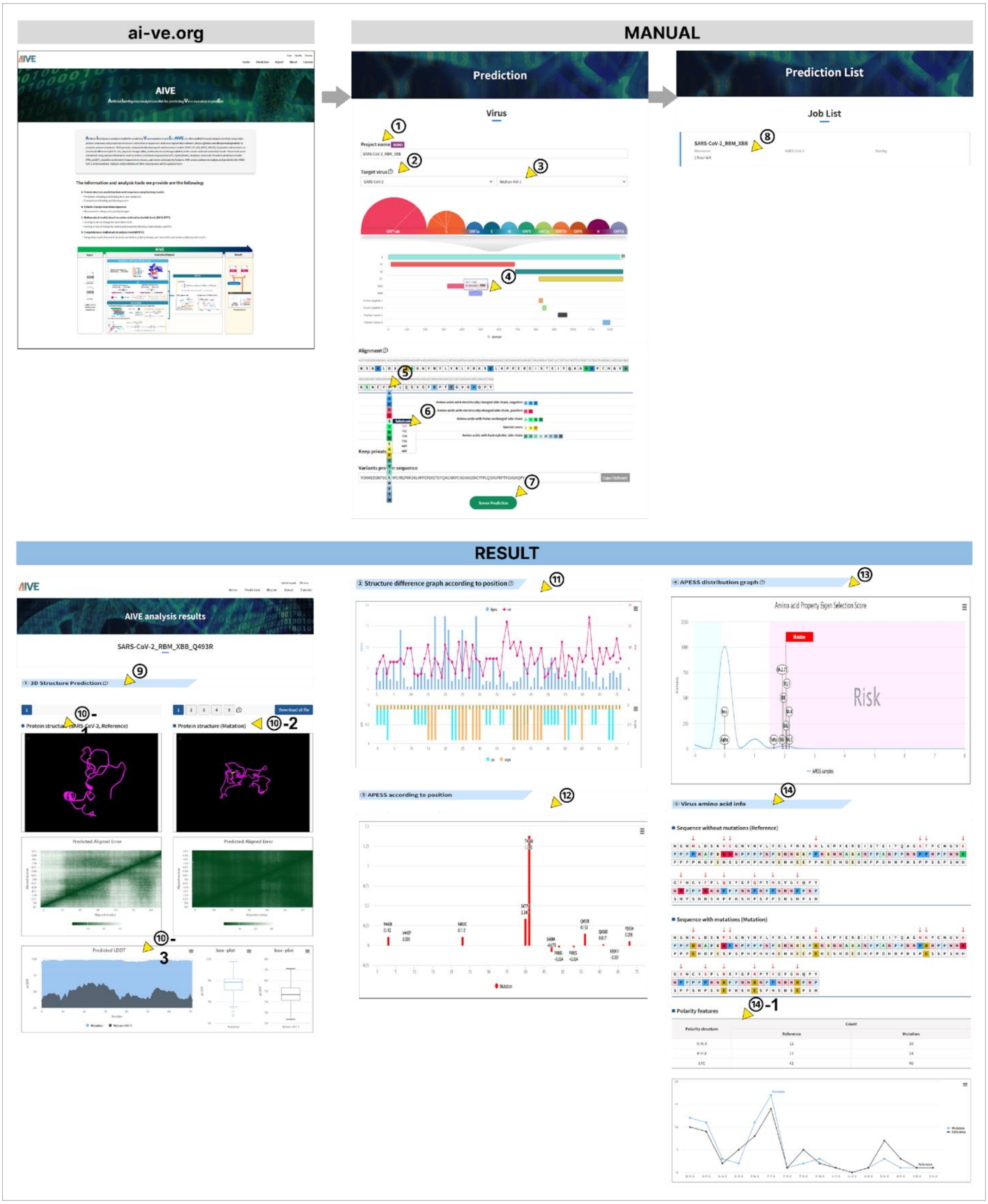

**Supplementary Figure 23. The user interface / user experience (UI/UX) of AIVE system.** An instruction manual for accessing the AIVE website (ai-ve.org) and using its prediction was shown. ① The project name is entered by user. ② The target virus is selected. In the demonstration, SARS-CoV-2 is selected. ③ The specific Variant of Concern (VOC) can be selected. In the demonstration, Wuhan-Hu-1 is selected. ④ Click on the RBD region. ⑤ The amino acid can be selected in the position in the sequence. ⑥ The codon can be selected for the amino acid. ⑦ Server prediction is started. ⑧ For Prediction List, it shows the current job list with ongoing projects and how much time is left for analysis. ⑨ The 3D Structure Prediction results are shown. **⑩-1** The 3D Protein structure (SARS-CoV-2, Reference) **⑩-2** The 3D Protein structure (Mutation). **⑩-3** The Predicted LDDT score. ⑪ Scores of SCPS, PCS, MR, BPES by amino acid position. ⑫ Scores of APESS by amino acid position. ⑬ Distribution of APESS score of SARS-CoV-2 VOCs Alpha, Beta, Delta, and Omicron (BA.2.75, BA.5, XBB, BQ.1) and location of APESS range as the input. ⑭ Comparison of polarity change between input and reference. **⑭-1**. Number of NN*, NP*, NA*, NB*, PN*,PP*, PA*, PB*, AN*,AP*, AA*, AB*, BN*, BP*, BA*, BB* between input and reference.

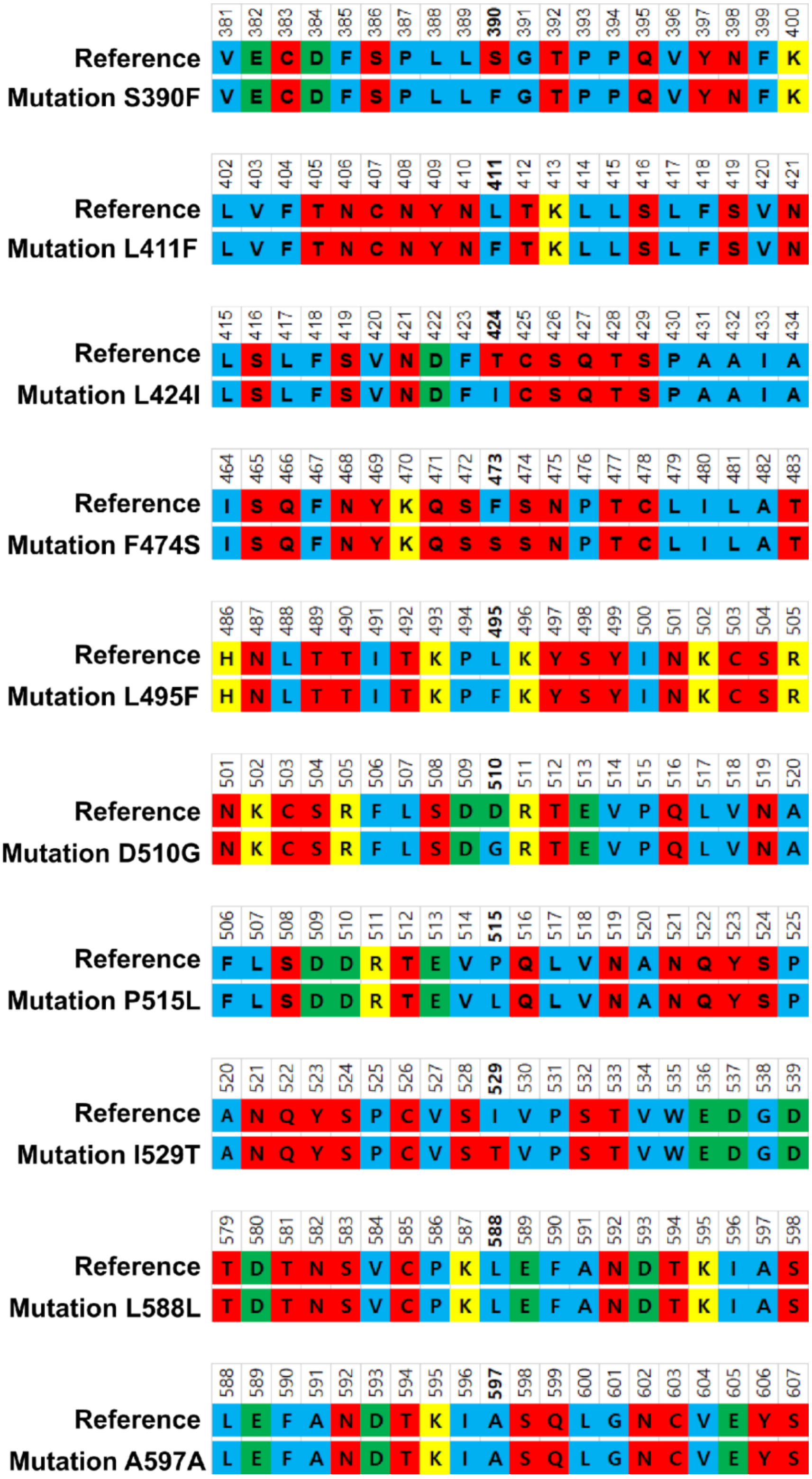

**Supplementary Figure 24. Polarity change analysis due to mutation in the amino acid sequence of MERS-CoV.** The amino acid sequence for MERS-CoV, a known coronavirus like SARS-CoV-2, and the mutations are displayed. The red indicates hydrophilic (polar), the blue indicates hydrophobic (non-polar), the green indicates acidic, and the yellow indicates alkaline (basic) amino acids. In MERS-CoV, a limited number of mutations were found. Notably, the S390F, L411F, T424I, F473S, D510S, P515L, I529T, L588L, and A597A mutations. In characteristics of MERS-CoV mutations, consecutive hydrophilic patterns are found in MERS-CoV, which the mutations do not disrupt.

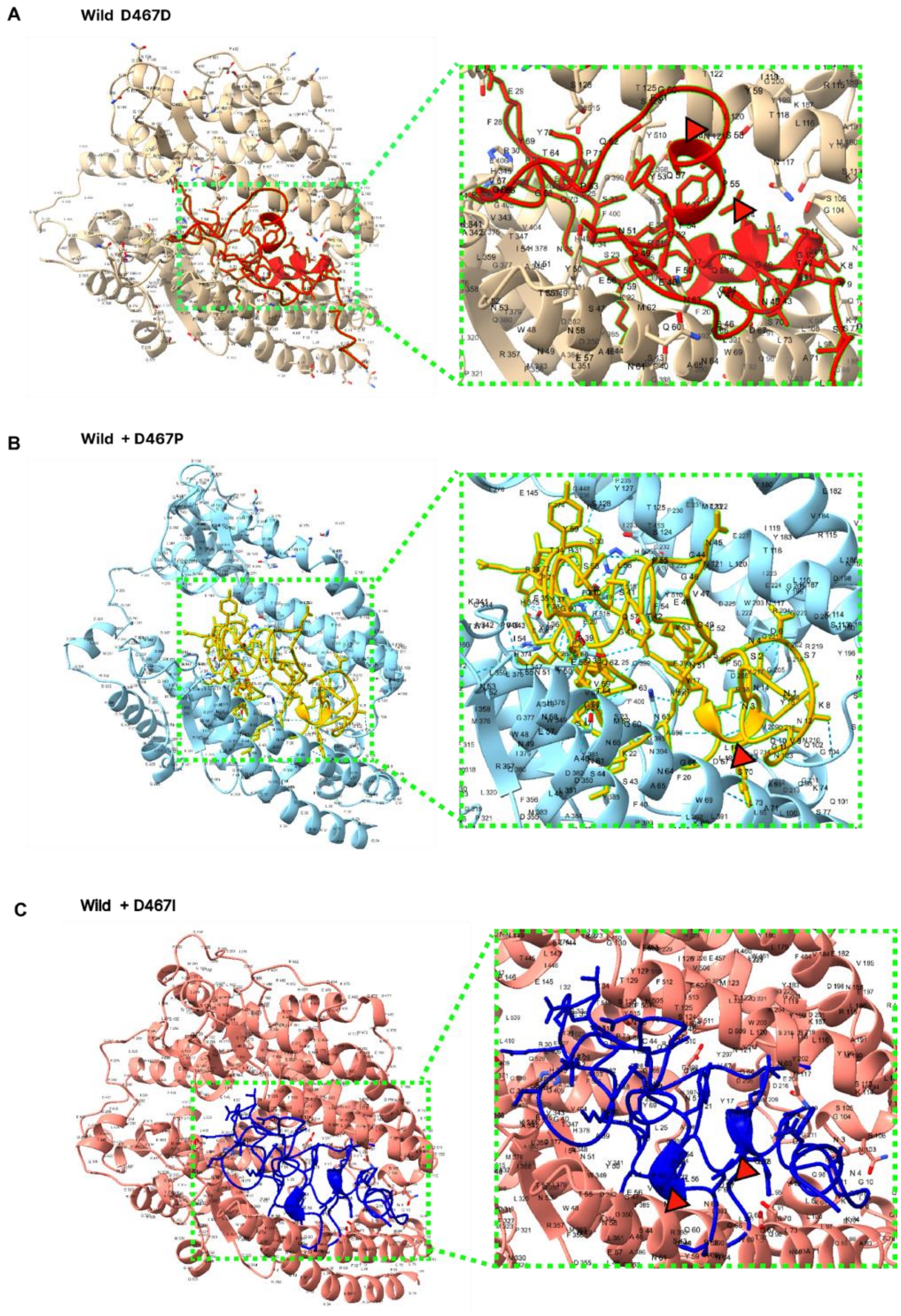

**Supplementary Figure 25. D467 protein 3D structure prediction.** Measuring the viral infectivity in vitro of the SARS-CoV-2 RBM sequence showed the structural importance of D467 (Figure 2C). In silico results of measuring the docking score between SARS-CoV-2 RBM and the host ACE2 receptor showed contradicting results (Figure 7A). Visualization of the SARS-CoV-2 folding structure displayed the wildtype D467D with an alpha-helix structure, indicated with a red triangle. Mutagenesis to D467P and D467I to the wildtype backbone resulted in the alpha helix structure turning into a linear structure. It becomes more likely for the protein 3D structure to become flexible.

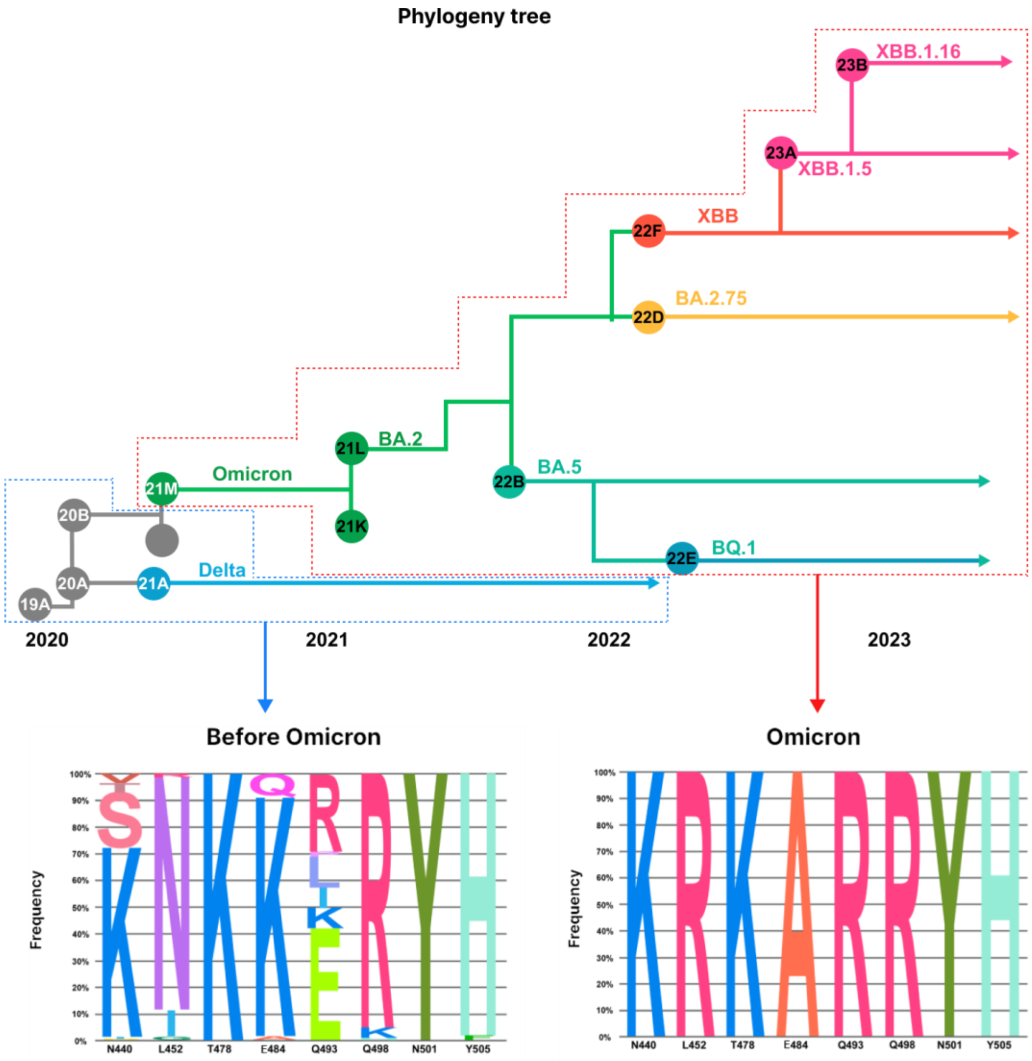

**Supplementary Figure 26. Frequencies of amino acid substitutions of major variants within clades.** Phylogenetic tree showing the evolution of SARS-CoV-2 lineages and their sub-lineages. Before Omicron, at various positions N440, L452, T478, T484, Q493, Q498, N501, Y505, the frequency of amino acids is displayed. There are different types of amino acid substitutions present with the different positions showing varied percentages. In Omicron lineages, the frequency of amino acid substitutions of N440K, L452R, T478K, T484A, Q493R, Q498R, N501Y, Y505H approach 100%, showing that amino acid substitutions become fixed.

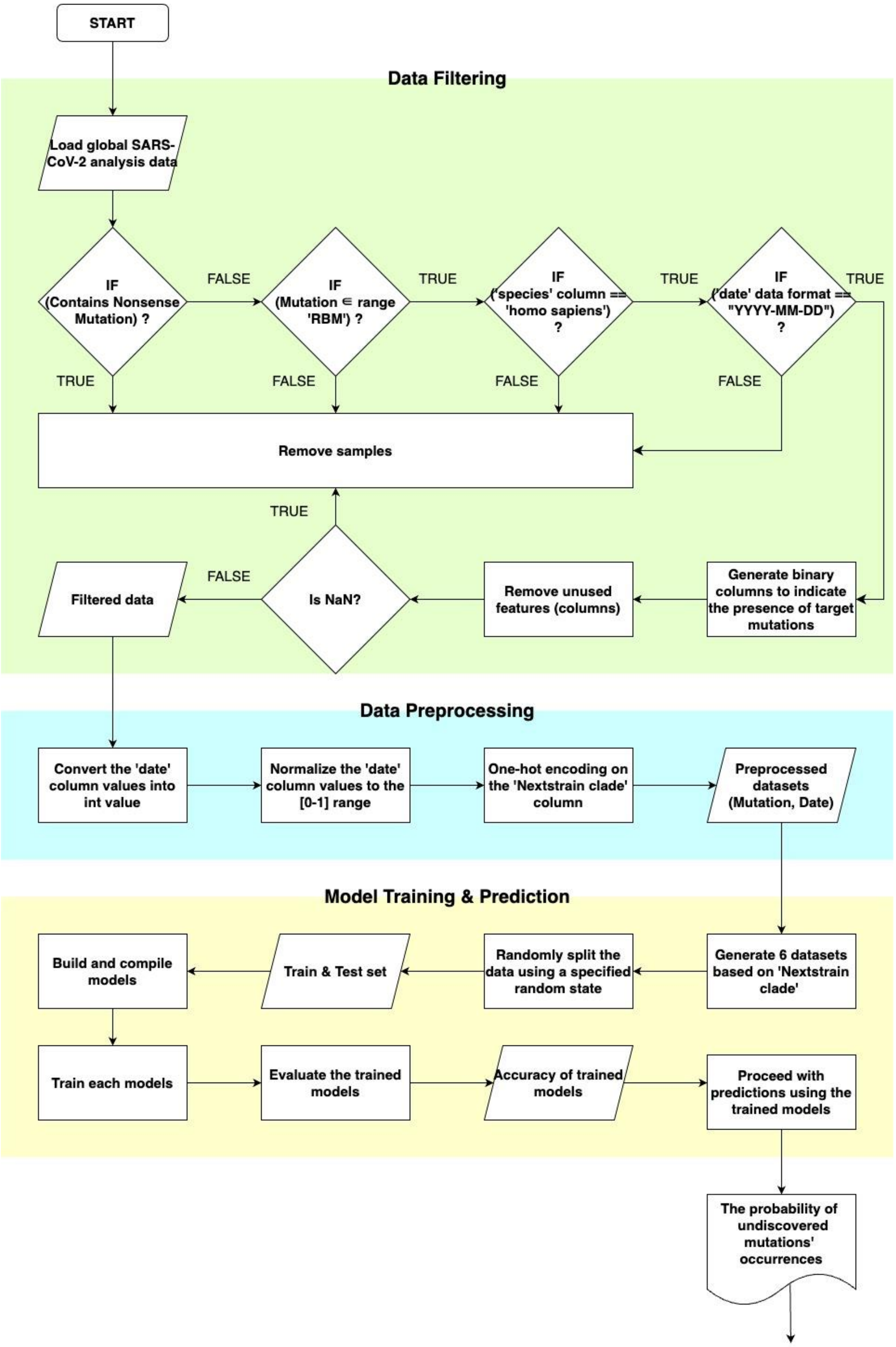

**Supplementary Figure 27. FlowChart for predicting mutation occurrences using artificial intelligence.** We used machine learning and deep learning methods to predict the likelihood of mutations occurring. ① We began by filtering the information and samples from the GISAID data. Unnecessary samples were removed depending on the presence of nonsense mutations, the host species, and the data format of the date column. Then, target mutations in the RBM region were converted into binary variables. Finally, filtering is completed by removing unused features and samples with missing data, excluding the clade, date, and binary target mutation columns. ② In order to utilize the filtered data for training, we preprocessed the data. We replaced the date information in the YYYY-MM-DD format with integer values indicating how many days had passed since the earliest recorded date, 2019-12-23. Then, the values were normalized to [0-1]. We then created preprocessed data by one-hot encoding the clade data. ③ We created a training model then used the trained model to predict the likelihood of the target mutations. Five different training models were involved in this process. ③-1 A prediction model using LightGBM, ③-2 a model using XGBoost, ③-3 sklearn library's Random Forest model, and then an ensemble model of the previous three models were used. Finally, ③-4 keras was used to create a multi-output neural network model for training and predicting the likelihood of the target mutations.

### Sign up & Login

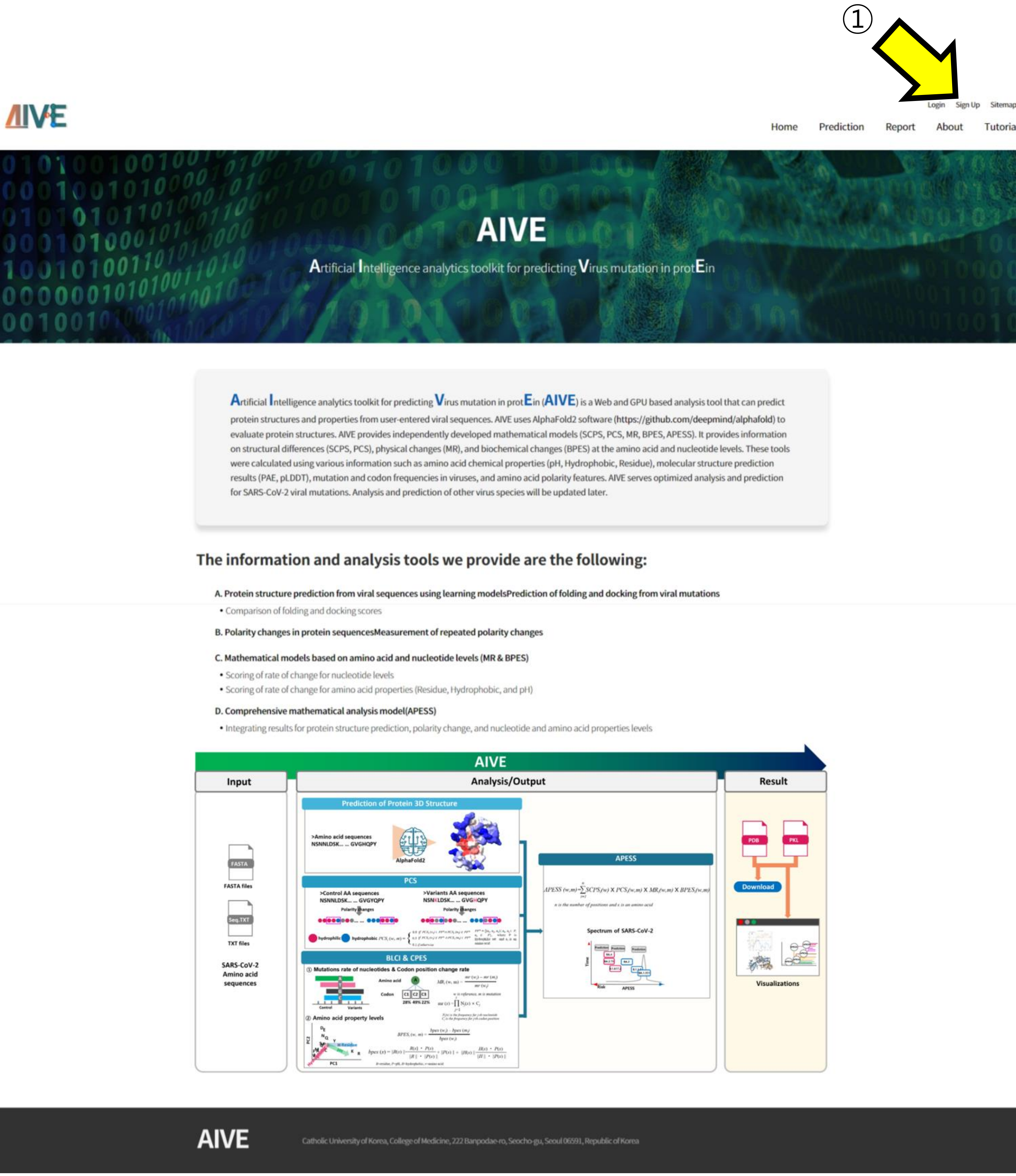

AIVE is used by creating a personal account for the ease of comparing analysis results.
① Click on the "Sign Up" option to proceed with the account creation.

### Sign up & Login

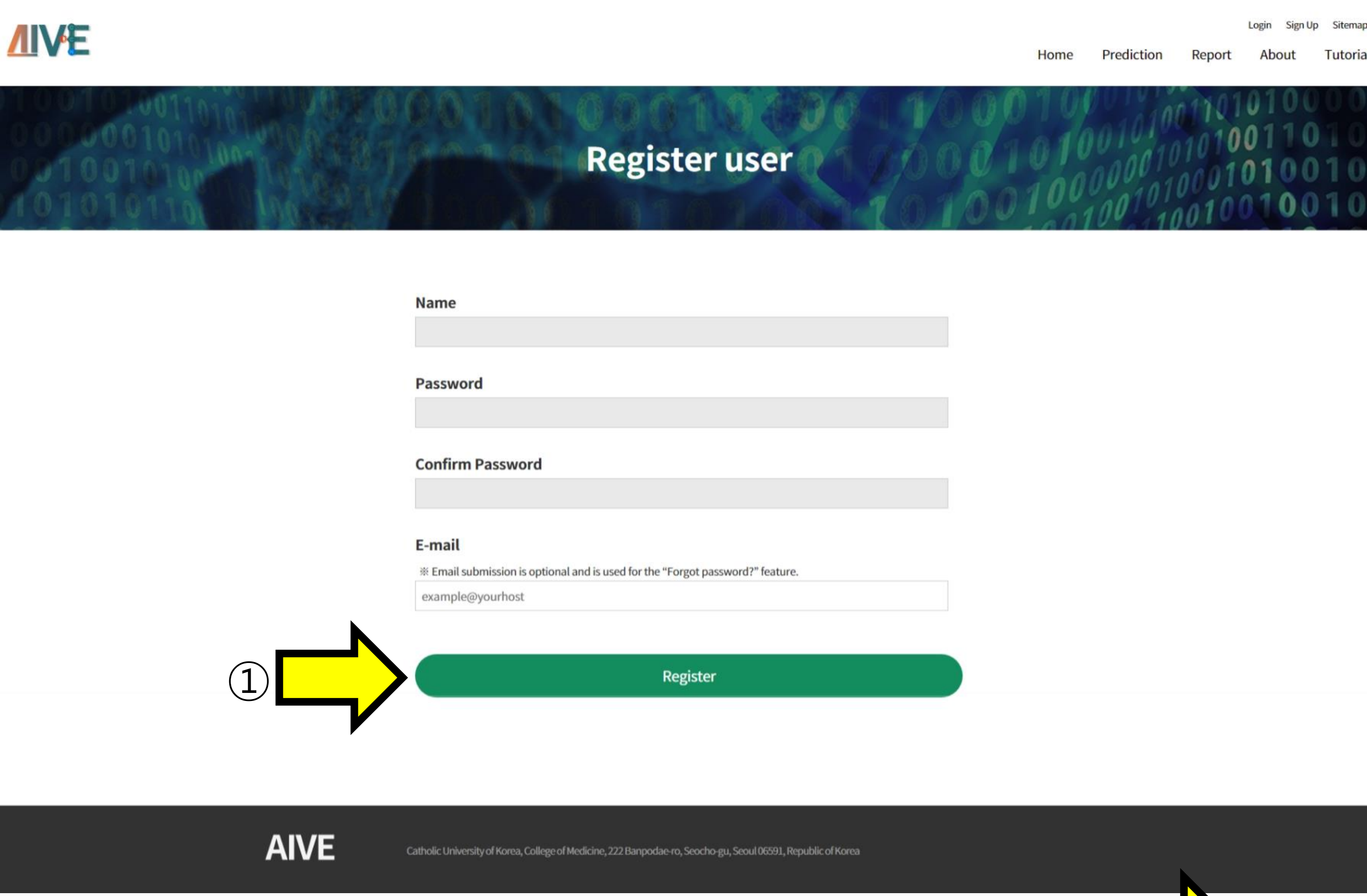

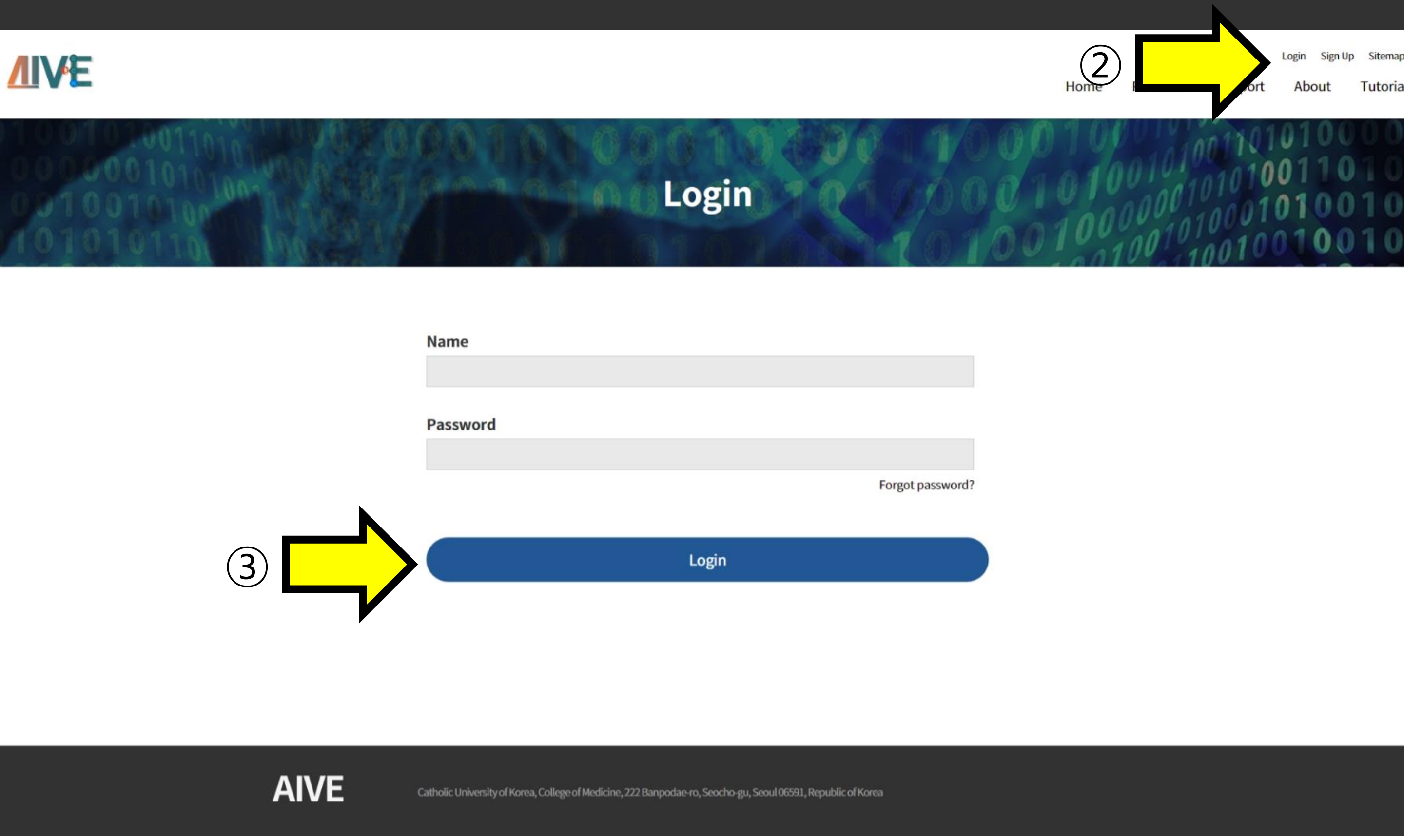

① Enter the username and password. To use the "Forgot password?" feature, enter your email and proceed with account creation.
②, ③ Access the "Login" section, enter the account information, and log in.

### Demo

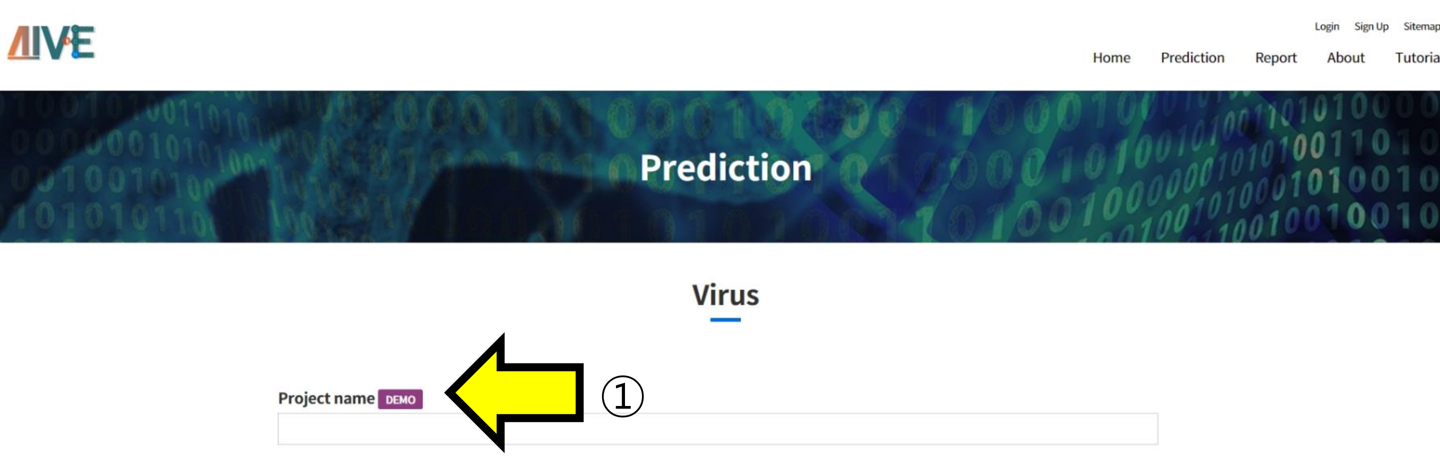

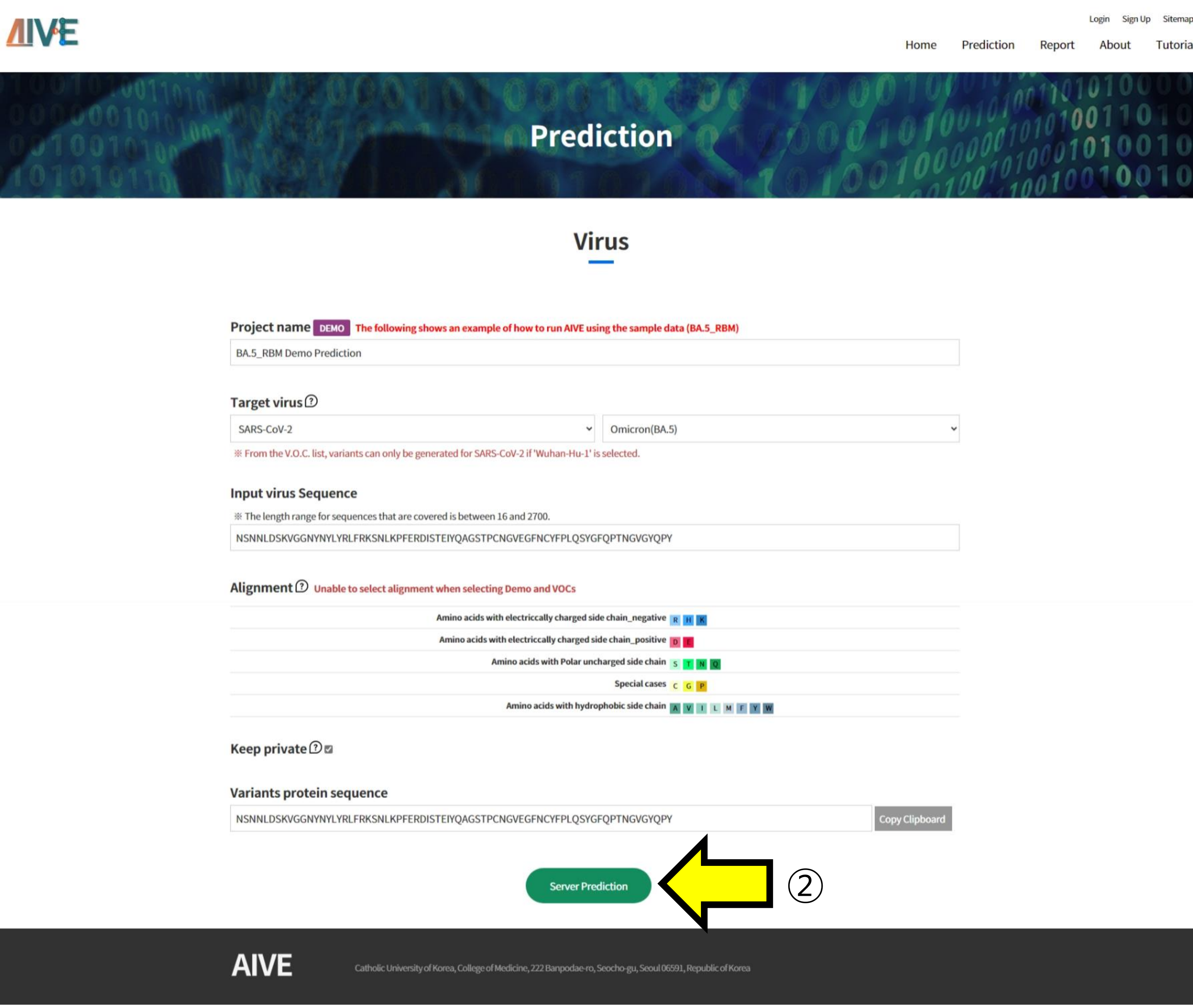

AIVE provides a demo feature that demonstrates the analysis of SARS-CoV-2 variants of concern (VOCs).
① Clicking on the Demo button will automatically select the RBM [S:437-508] region of the SARS-CoV-2 BA.5 variant.
② To predict and analyze the structure of the sequence generated by the Demo function, click the "Server Prediction" button to submit the task.

### Predict VOC structure

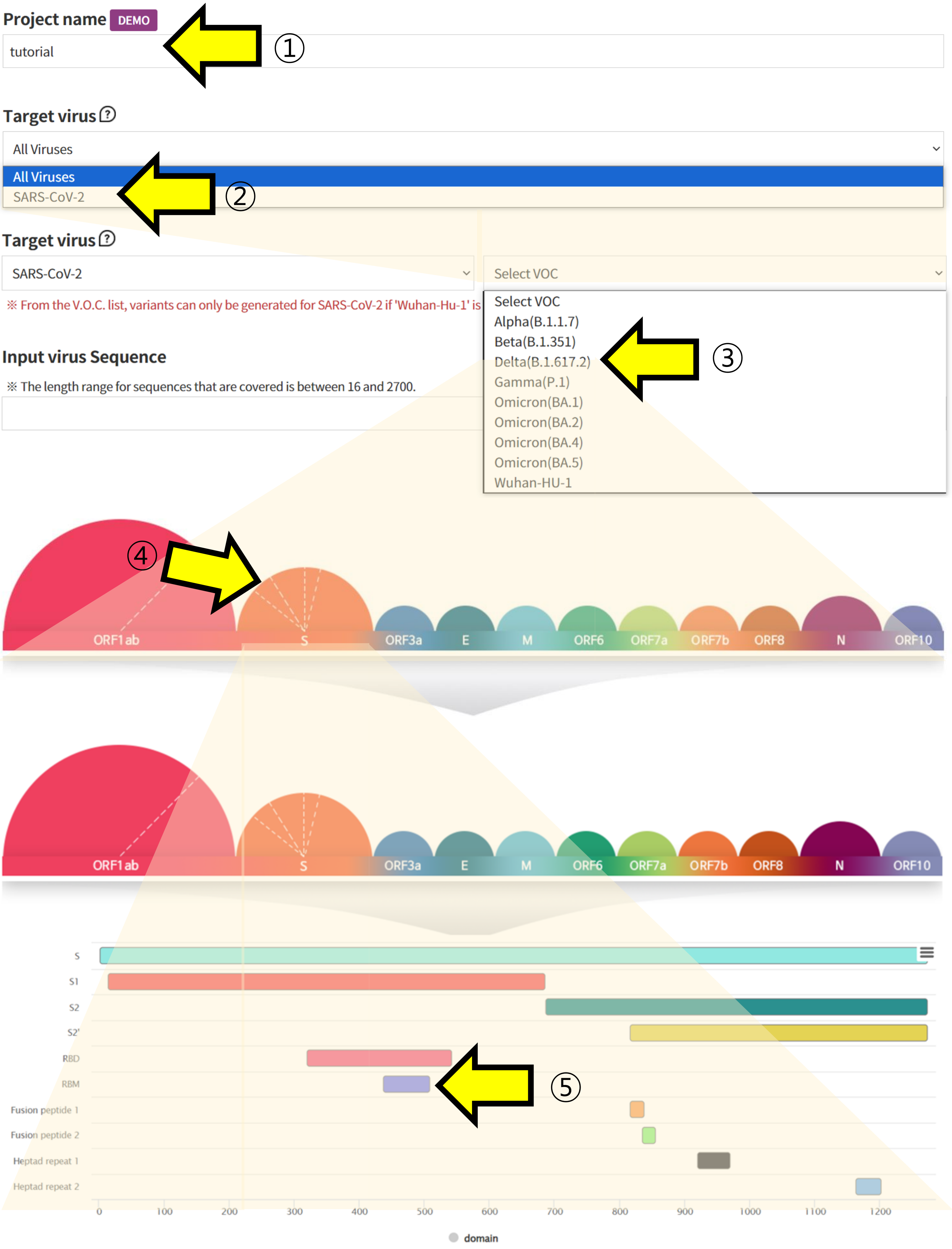

In AIVE, users can personally select and analyze the VOC and domains of SARS-CoV-2.

① Write the Project name for task categorization.

② Select SARS-CoV-2 in the Target Virus section to retrieve information on Coronaviruses.

③, ④, ⑤ Choose the VOC and regions you want to analyze to retrieve the corresponding Amino Acid sequence information.

### Submit

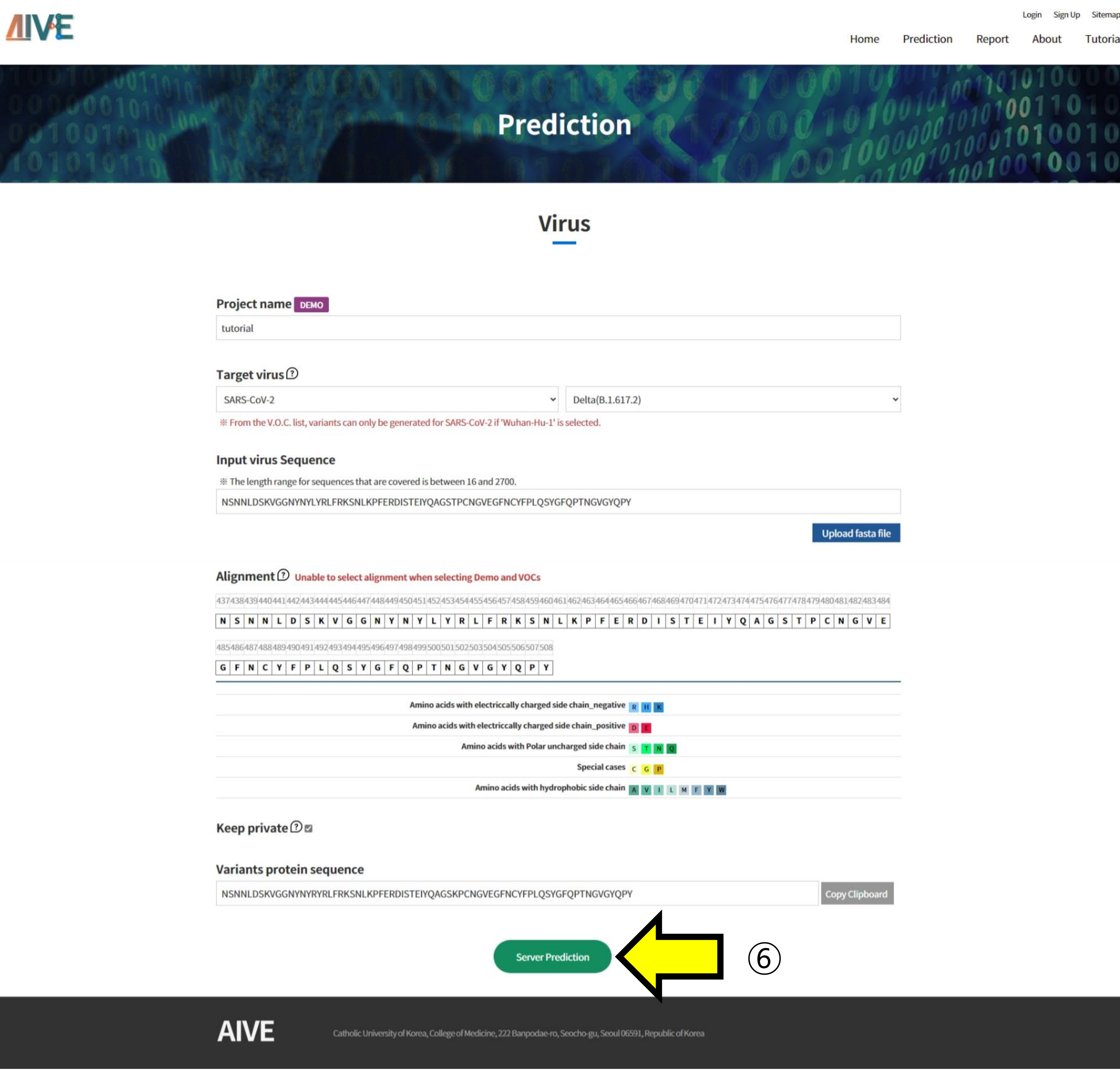

⑥ Click the "Server Prediction" button to start the prediction and analysis of the Project

### List

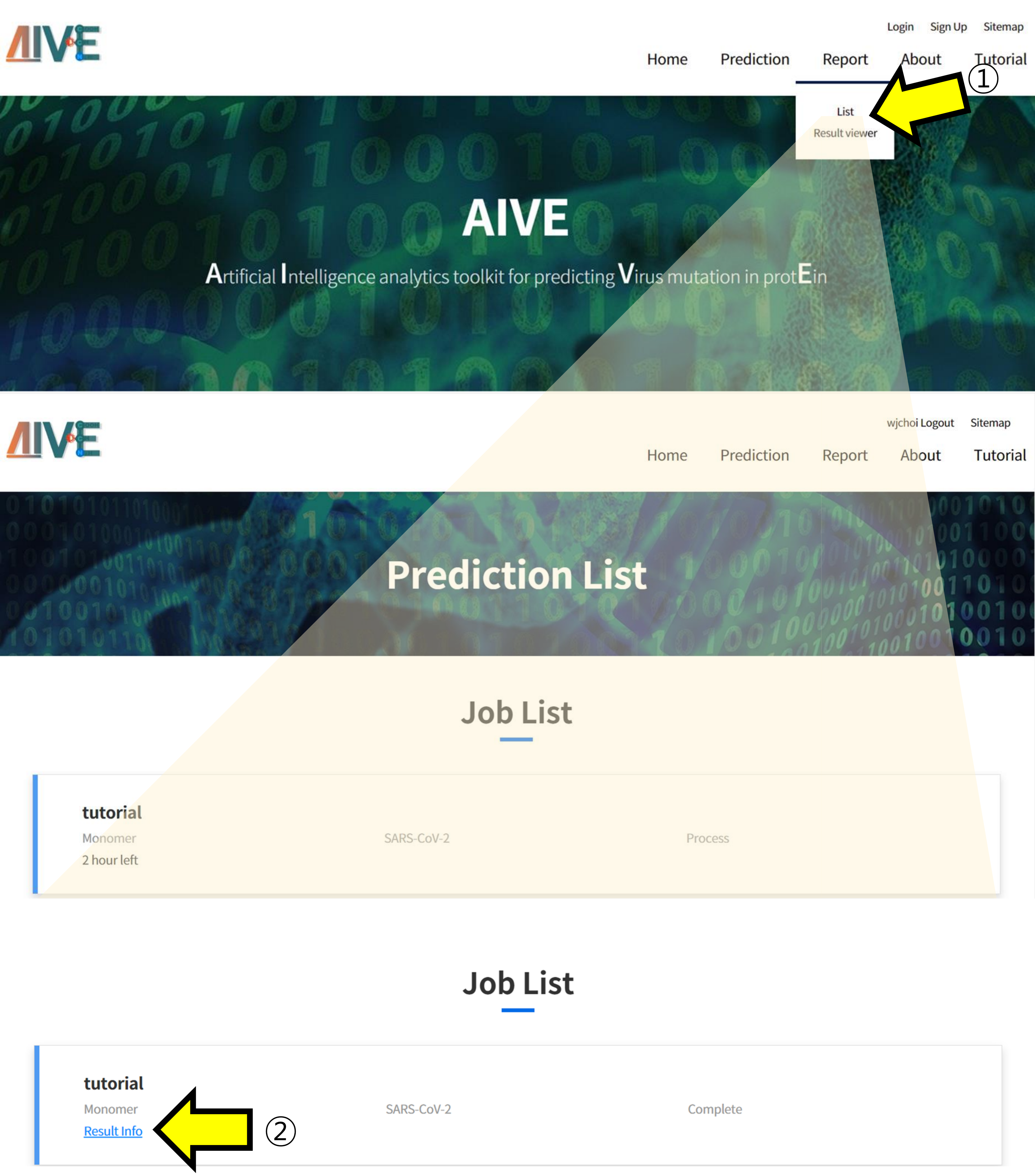

In the "List" section, users can check the list of submitted Projects and their progress.
① Click the "Report menu" and then select "List" from the submenu to check Projects.
② You can check the analysis results by clicking on the "Result info" section for completed tasks.

### Generate SARS-CoV-2 mutated sequence

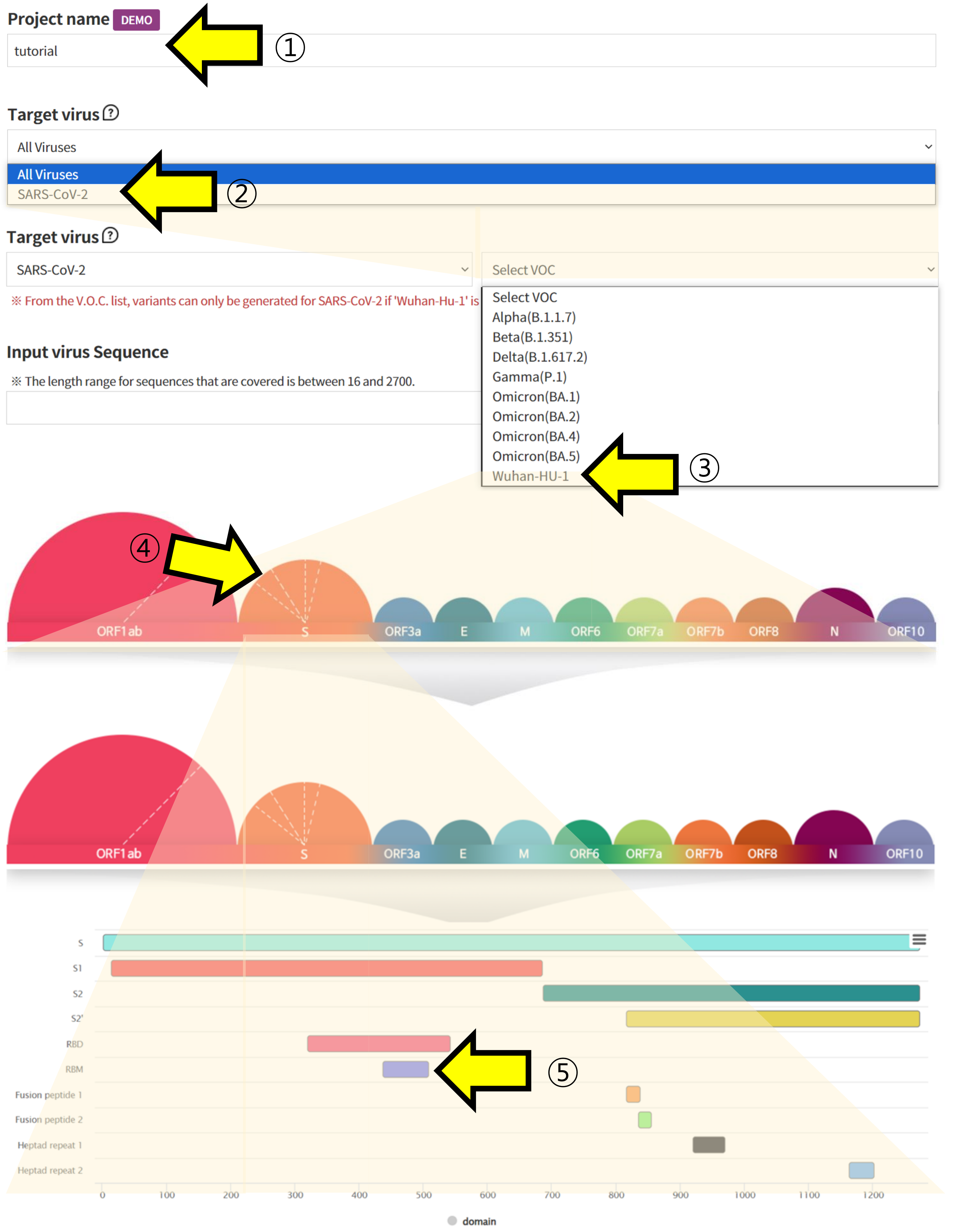

Users can not only access information about VOCs but also generate mutations for analysis.

① Specify a Project name to categorize the submitted task

②, ③, ④, ⑤ Choose Wuhan-HU-1 sequence of SARS-CoV-2 and retrieve the sequence of the region you want to verify.

### Generate SARS-CoV-2 mutated sequence

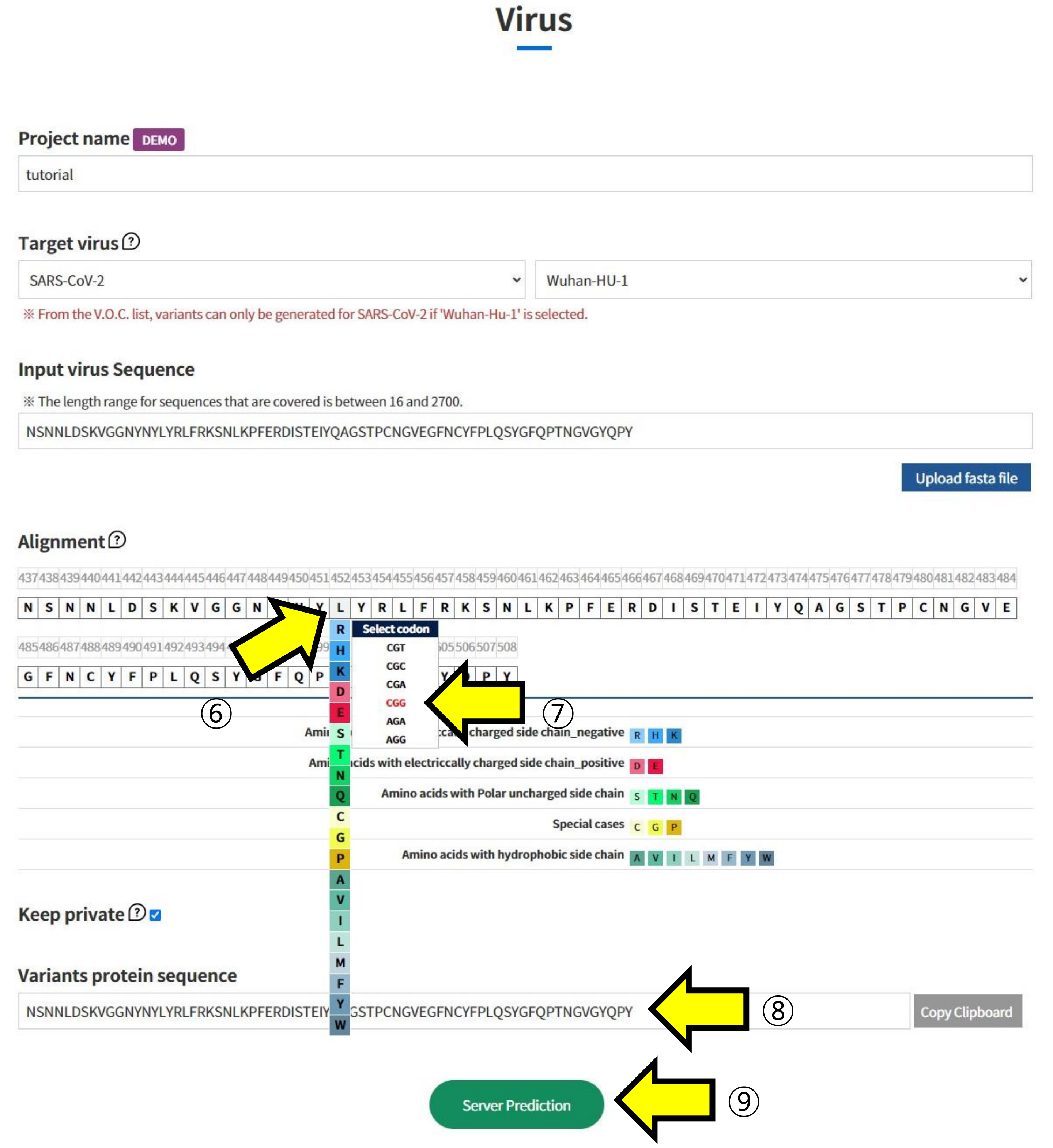

⑥ In Alignment, click on the positions in the sequence of Wuhan-HU-1 to select the mutated Amino Acid.
⑦ Additionally, choose which codon to mutate into from the selected Amino Acid. This function is used to assess the impact of mutations at the codon level.
⑧ The sequence with the selected mutations is displayed.
⑨ Submit the task to proceed with the structural prediction and analysis of the mutated sequence.

### User sequence - Monomer

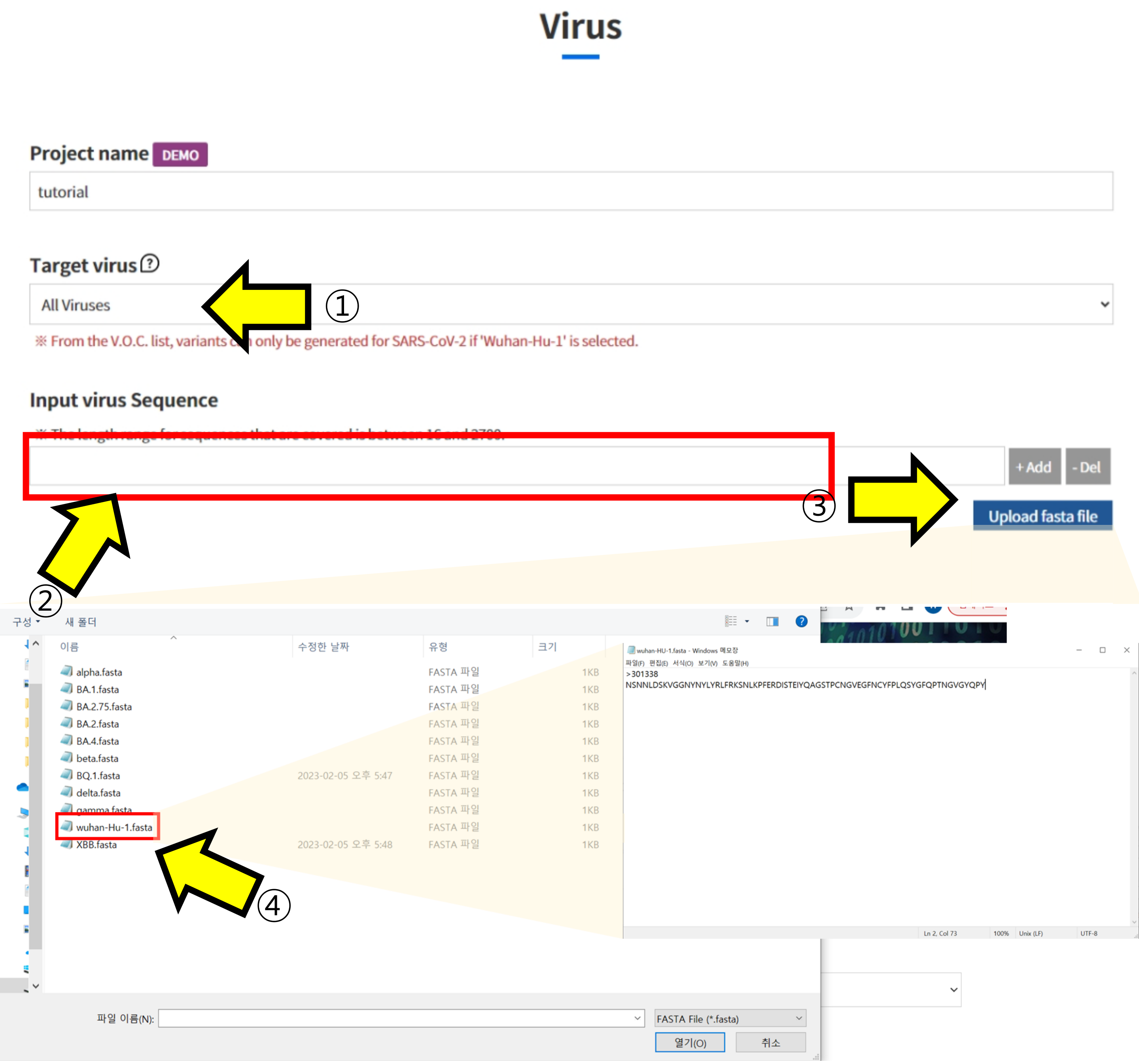

In AIVE, users can predict the structure of sequences, not limited to SARS-CoV-2. Let's take a look at monomer structure prediction

① Select "All viruses" in the Target Virus section.

② Enter the Amino Acid sequence you want to check in the "Input virus Sequence" box.

③ Alternatively, you can upload a fasta file instead of entering it directly.

④ When the upload window appears, select the fasta file you want to check to pull up the sequence.

### User sequence

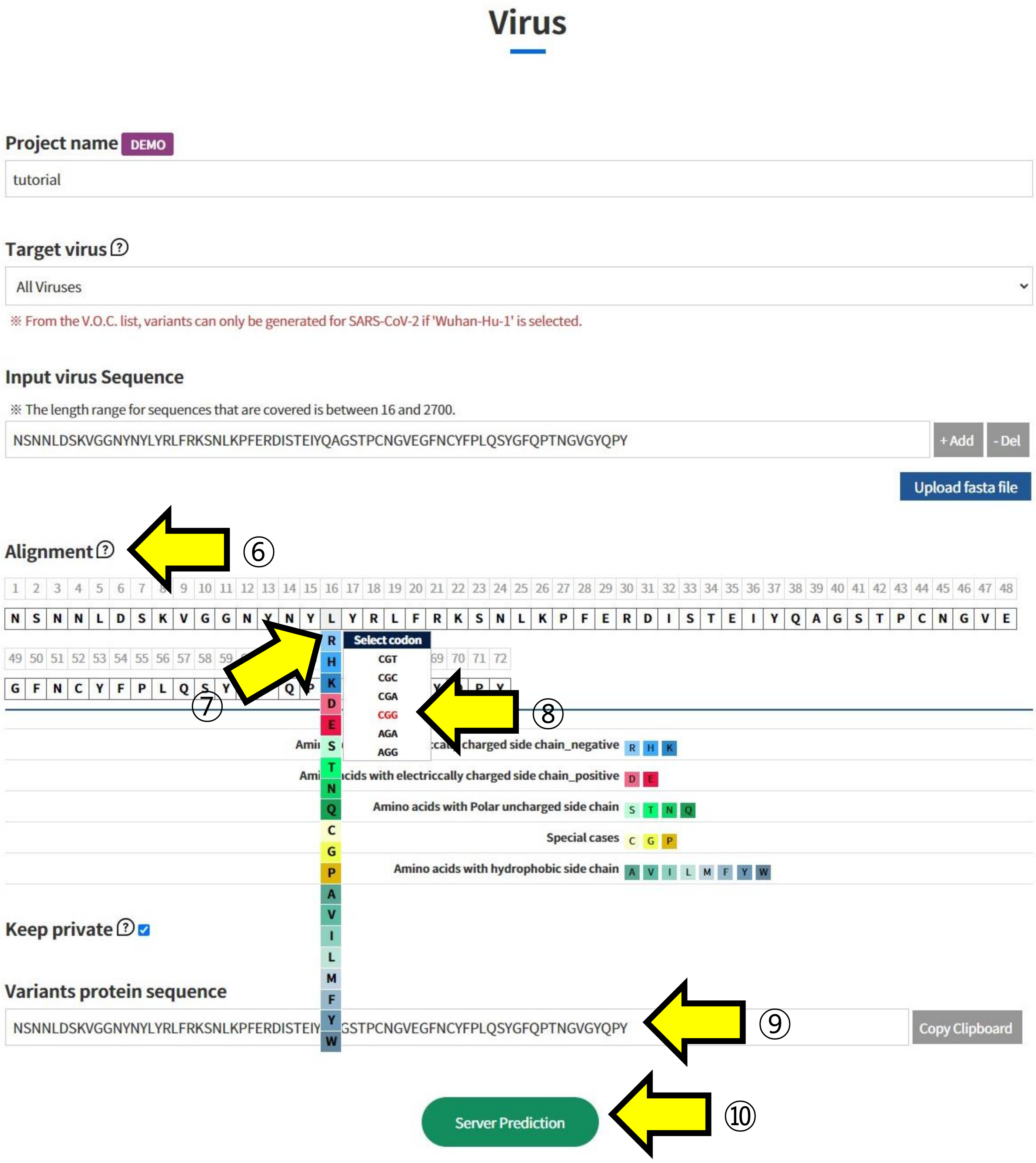

⑥ The Amino Acid sequence recorded in the fasta file you uploaded is displayed in “Alignment”.
⑦ Click on the positions in the displayed sequence where you want to introduce mutations and select the mutated Amino Acid.
⑧ Additionally, choose which codon to mutate into from the selected Amino Acid. This function is used to analyze the impact of mutations at the codon level.
⑨ The sequence with the selected mutations is displayed.
⑩ Submit the task to proceed with the structural prediction and analysis of the mutated sequence.

### User sequence - Multimer

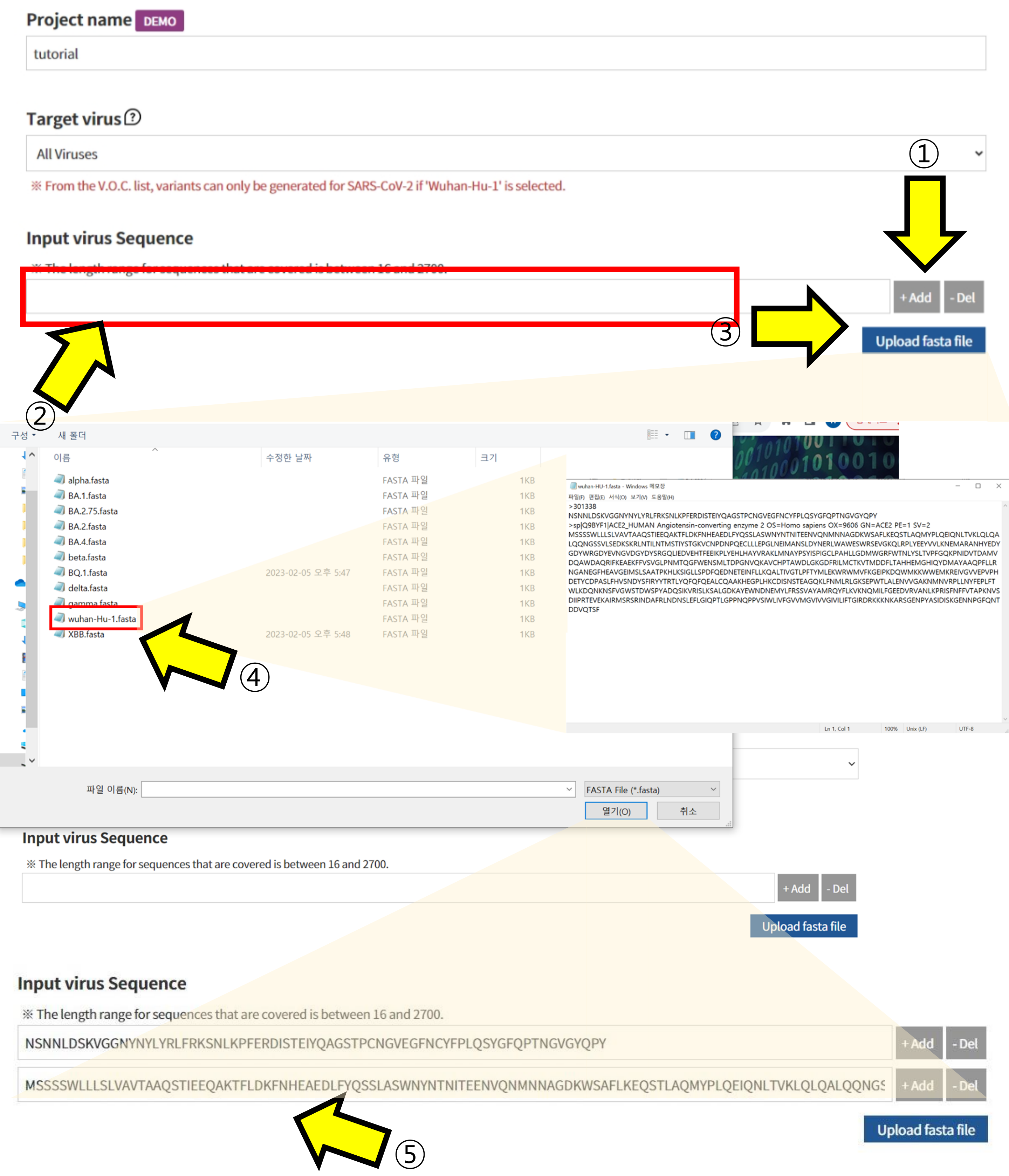

Let's look at the case of predicting a Protein Complex structure:

① Click the +Add button to create as many sequence input boxes as there are chains in the protein complex you want to predict.

② Enter the Amino Acid sequence of each chain in the generated “Input virus Sequence” boxes.

③ Alternatively, a fasta file can be uploaded without entering the sequence directly

④ When the upload window appears, select the fasta file to retrieve the sequence.

⑤ The sequences of each chain, as stored in the fasta file, are inputted.

### User sequence - Multimer

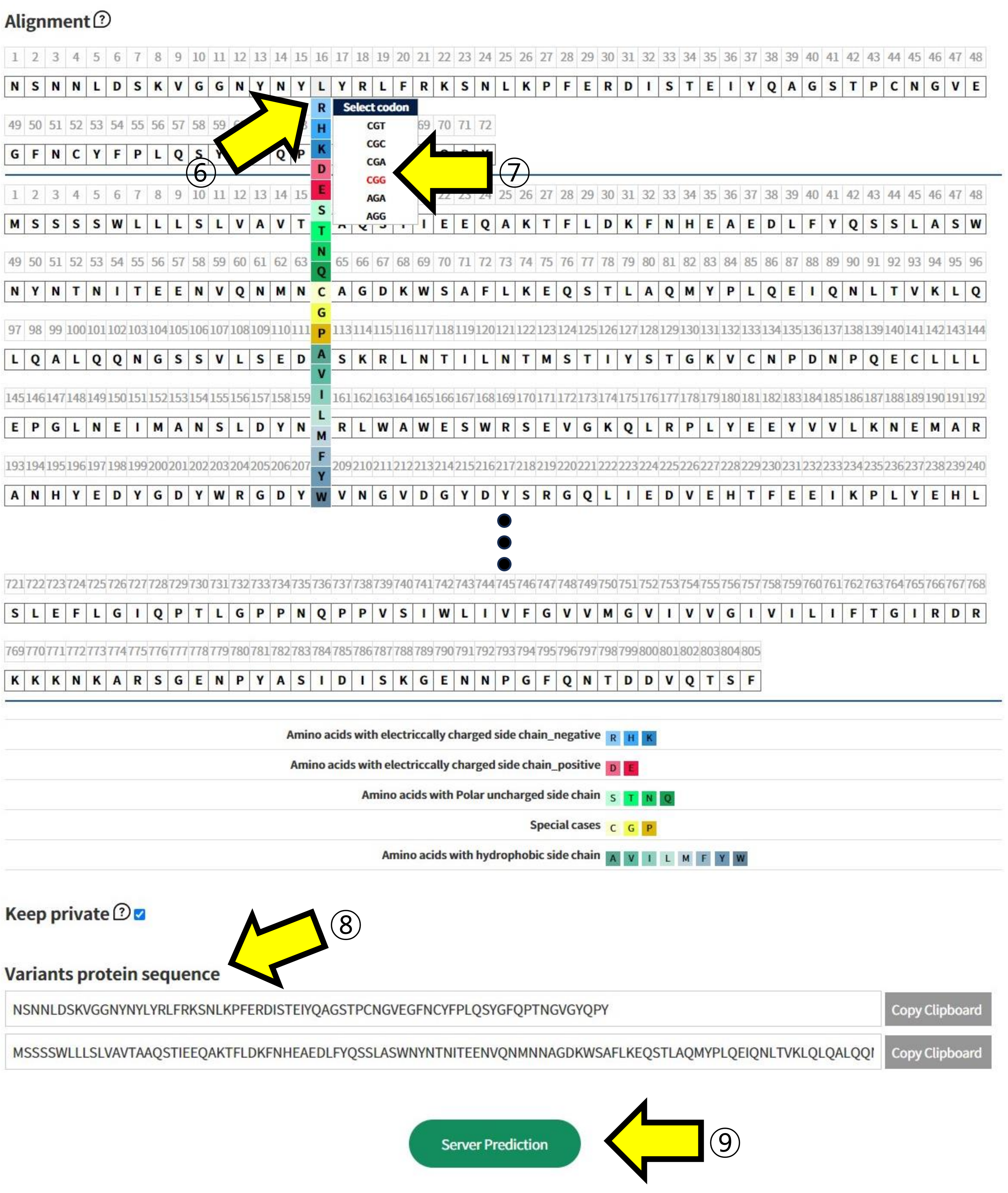

⑥ Click on the positions in the displayed sequences where you want to introduce mutations and select the mutated Amino Acid.
⑦ Additionally, choose which codon to mutate into from the selected Amino Acid. This function is used to analyze the impact of mutations at the codon level.
⑧ The sequence with the selected mutations is displayed.
⑨ Submit the task to proceed with the structural prediction and analysis of the mutated sequence.

### Result report page - Structure

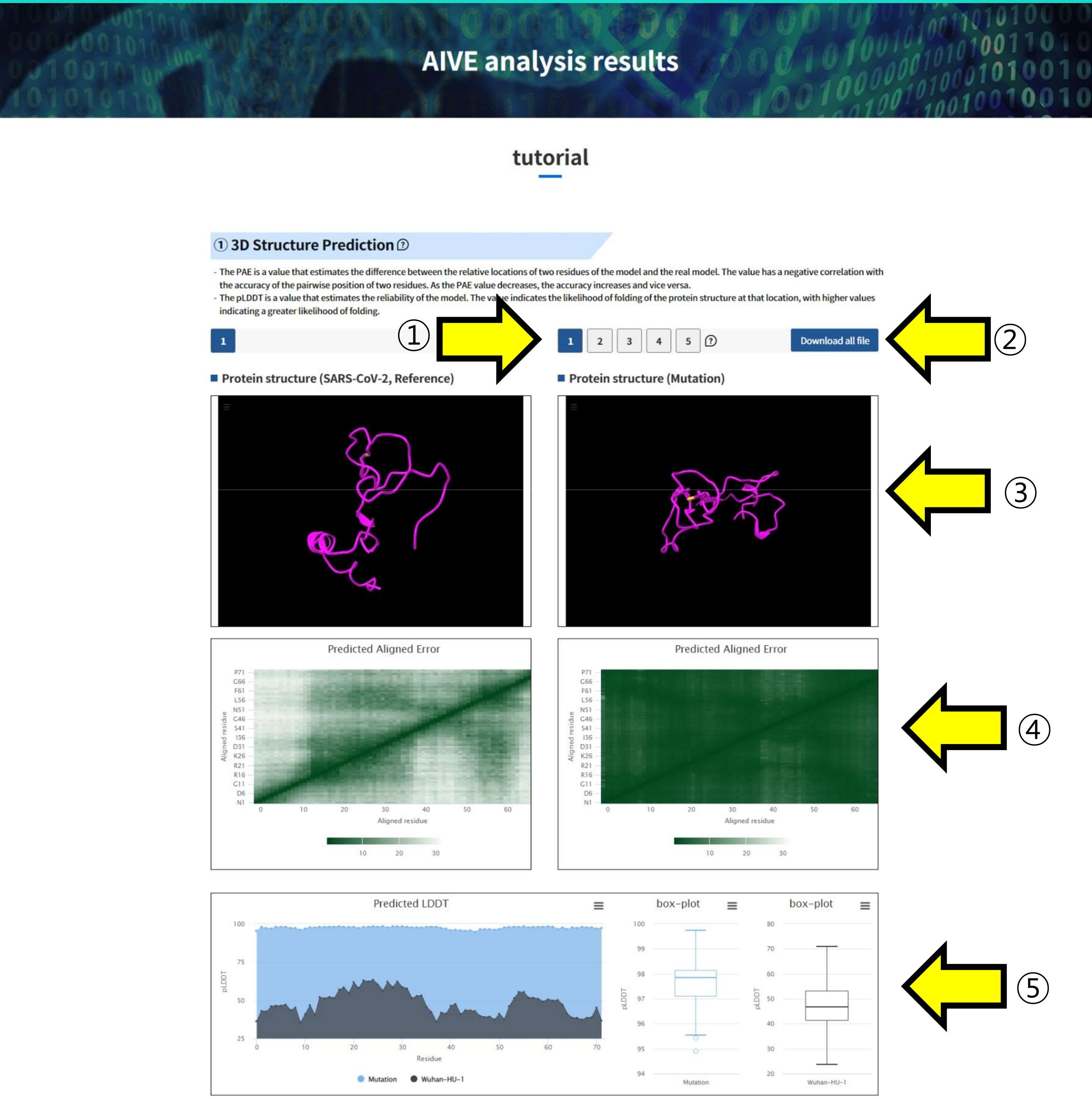

From the prediction and analysis results of SARS-CoV-2, the 3D structure prediction results can be accessed.

① You can select and view the predicted 3D structures by choosing from the five available options.

② Use the "Download all file" button to download the result files of the predicted structures to your device.

③ You can visualize the predicted 3D structure for inspection and comparison with SARS-CoV-2 Wuhan-HU-1. ④, ⑤ Clicking on the highlighted regions in positions allows you to inspect them.

④ Predicted aligned error (PAE) is a value that estimates the difference between the relative locations of two residues of the model and the real model. A low PAE value indicates that the accuracy of the relative location of the two residues is high.

• The color at (x, y) indicates AlphaFold's expected position error at residue x if the predicted and true structures were aligned on residue y.

• If the PAE is generally low for residue pairs x, y from two different domains, it indicates that AlphaFold predicts well-defined relative positions and orientations for them. (Explanation from AlphaFold FAQ)

⑤ Predicted LDDT(pLDDT) is a value that estimates the reliability of the model. It estimates how well the actual model residue and predicted model residue match. At the same time, it indicates how well the protein structure folds in the corresponding location. A low pLDDT value indicates that the reliability of the corresponding position is low and that it possesses a disordered structure.

### Result report page - APESS

#### ② Structure difference graph according to position ⓘ

- Protein structure prediction characteristics (SCPS: SubClustering of Protein Structure in ①) and polarity (PCS: Polarity Change Score) are shown.
- SCPS divides the residues constituting the 3D protein structure into several groups using K-means clustering. The indicated groups are those with a high proportion of WHO name variants.
- PCS assigns weight to a position having a specific structure (where P appears consecutively) by polarity features of amino acid.
- Ratio change in frequencies of amino acids sequences (mr: mutation rate) and Biochemical properties of amino acid sequencies (bpes: biochemical properties eigen score) are shown.
- MR is the rate of change of Amino Acid and its constituent nucleotides; the higher the rate, the higher the rate of change.
- BPES measures changes in the biochemical properties of amino acids at the site of a mutation. A high rate of change results in a high BPES value. Biochemical properties are measured by integrating amino acid residue, pH, and hydrophobic information.

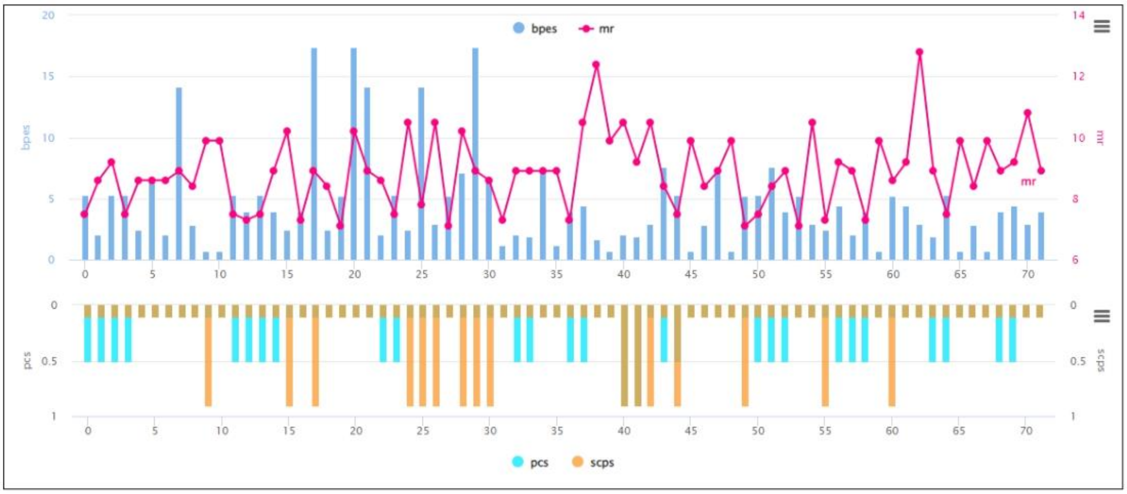

#### ③ APESS according to position

- APESS is an amino acid property eigen selection score calculated by multiplying the values of SCPS, PCS, MR, and BPES.
- APESS is a value that integrates the results of ② and ③. The higher the APESS value, the more dangerous the mutation is.
- The graph shows the positions of mutations with structural differences (SCPS, PCS) in viral proteins. It also shows the positions and magnitudes of relatively large physical changes (MR) and biochemical changes (BPES).

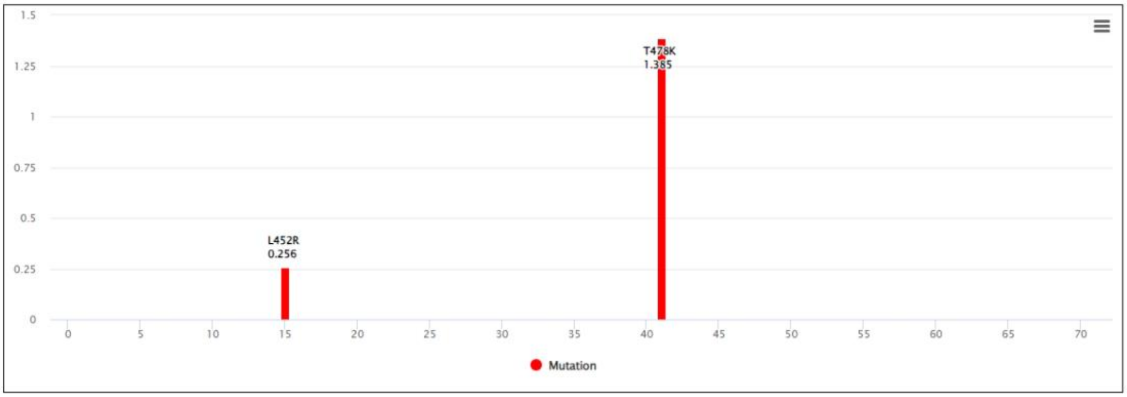

#### ④ APESS distribution graph ⓘ

- The following graph shows the APESS distribution of mutations generated through random sampling. The red area belongs to the quantile set of 0.05, signifying that the mutation is risky if it belongs to this area. A lettered balloon indicates the score position of the WHO VOC variant, and the red flag indicates the APESS position of the protein predicted in ①.

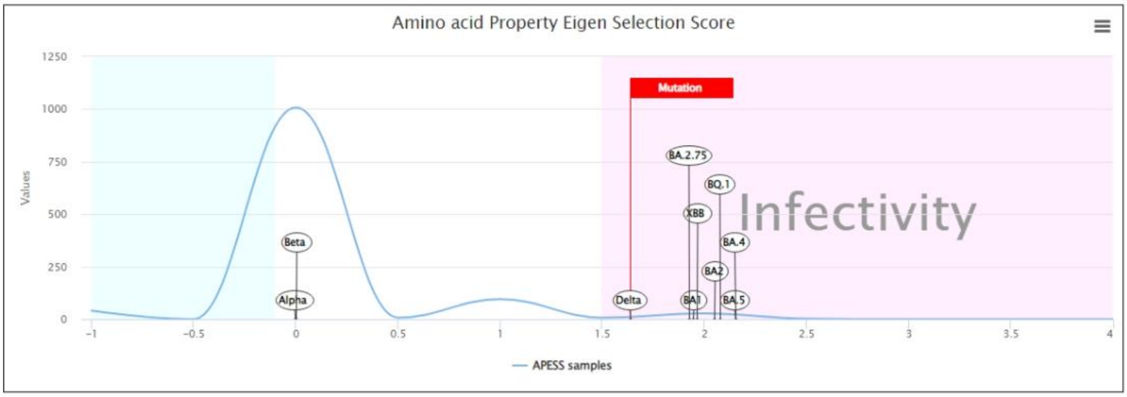

### Result report page – APESS subscore

#### ① SCPS

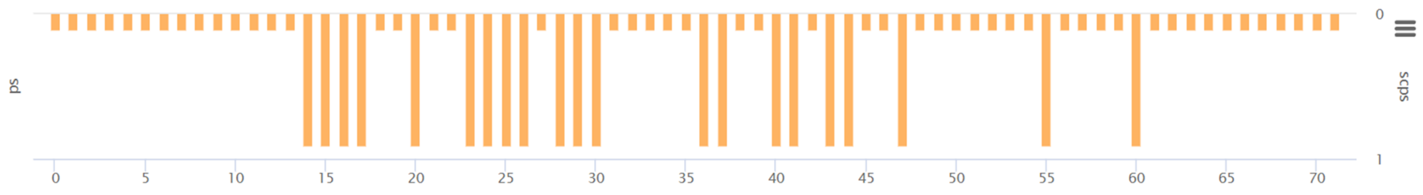

#### ② PCS

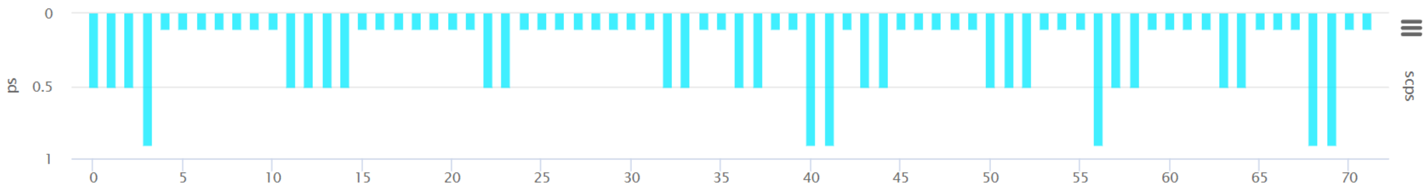

#### ③ BPES

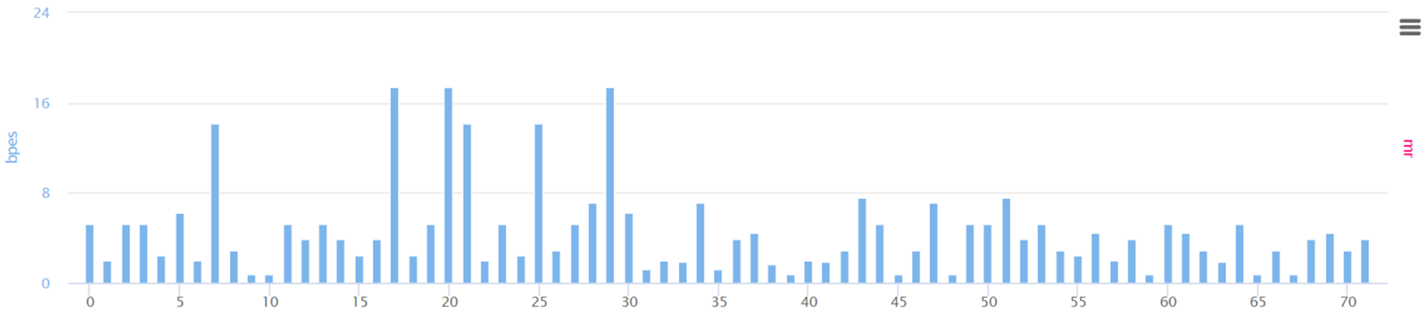

## ④ MR

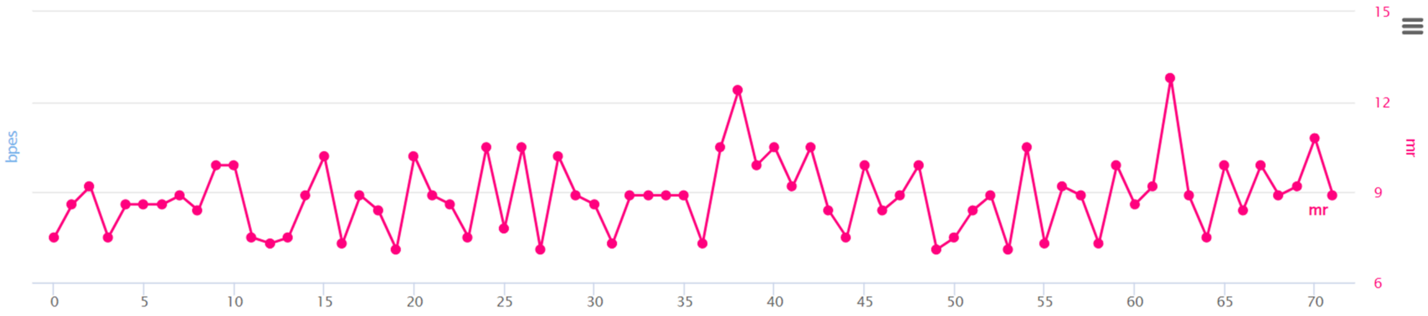

For the amino acid sequence entered by the user, the AIVE system provides a total of 6 evaluation charts.
① AIVE predicts protein structures due to mutations in each gene for coronavirus lineages or sub-lineages. From the predicted result, it carries out grouping of amino acids (components of 3D protein structure) utilizing K-means clustering to report SCPS results.
② through repeated pattern analysis of polar amino acids in amino acid sequences, AIVE reports PCS results.
③ AIVE figures out amino acid properties to measure BPES through measurement of changes in biochemical properties of amino acid.
④ AIVE calculates MR through rate of change for nucleotide frequencies due to mutations.

### Result report page – APESS & distribution

#### ⑤ APESS

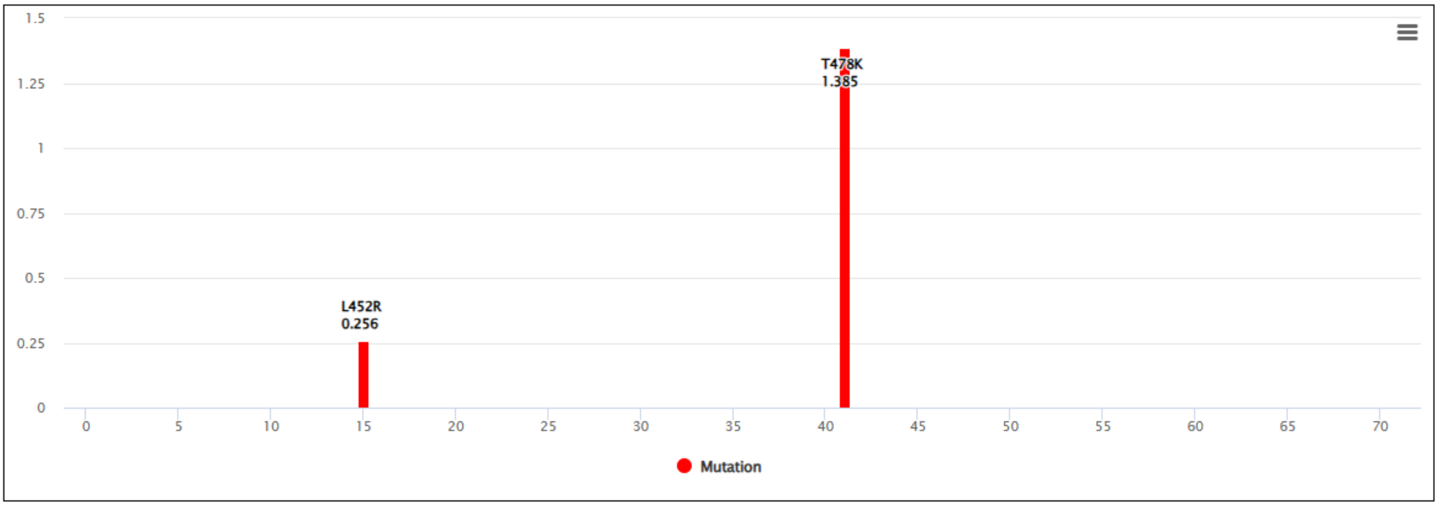

#### ⑥ APESS distribution graph

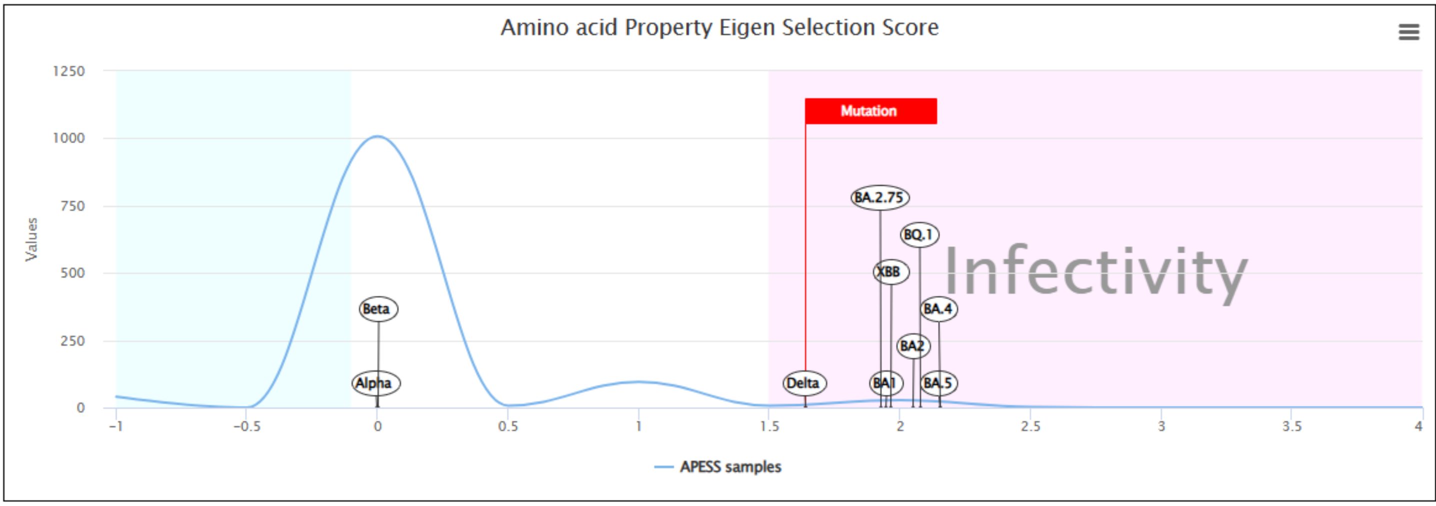

⑤ APESS, the result of comprehensive mathematical model (SCP*PCS*MR*BPES) of measured analysis results is provided.
⑥ the APESS distribution graph provides risk and spread results of the amino acid sequence entered by the user by comparing to VOCs' APESS evaluation metric.

### Result report page - Polarity

#### ⑤ Virus amino acid info

- Visualizes and tabulates changes in repeated polarity structure sequence for wild-type sequences and mutated type sequences used in ①.

##### Sequence without mutations (Reference)

| | | | | | | | | | | | | | | | | | | | | | | | |
|---|---|---|---|---|---|---|---|---|---|---|---|---|---|---|---|---|---|---|---|---|---|---|---|
| N | S | N | N | L | D | S | K | V | G | G | N | Y | N | Y | L | Y | R | L | F | R | K | S | N |
| P | P | P | P | N | A | P | B | N | N | N | P | P | P | P | N | P | B | N | N | B | B | P | P |
| P | P | P | P | H | O | P | E | H | S | S | P | H | P | H | H | H | E | H | H | E | E | P | P |

| | | | | | | | | | | | | | | | | | | | | | | | |
|---|---|---|---|---|---|---|---|---|---|---|---|---|---|---|---|---|---|---|---|---|---|---|---|
| L | K | P | F | E | R | D | I | [illegible] | [illegible] | [illegible] | I | Y | Q | A | G | S | T | P | C | N | G | V | E |
| N | B | N | N | A | B | A | N | P | [illegible] | [illegible] | N | P | P | N | N | P | P | N | P | P | N | N | A |
| H | E | S | H | O | E | O | [illegible] | [illegible] | [illegible] | O | H | H | P | [illegible] | [illegible] | P | P | S | S | P | S | H | O |

①

②

③

| | | | | | | | | | | | | | | | | | | | | | | | |
|---|---|---|---|---|---|---|---|---|---|---|---|---|---|---|---|---|---|---|---|---|---|---|---|
| G | F | N | C | Y | F | P | L | Q | S | Y | G | F | Q | P | T | N | G | V | G | Y | Q | P | Y |
| N | N | P | P | P | N | N | N | P | P | P | N | N | P | N | P | P | N | N | N | P | P | N | P |
| S | H | P | S | H | H | S | H | P | P | H | S | H | P | S | P | P | S | H | S | H | P | S | H |

④

##### Sequence with mutations (Mutation)

| | | | | | | | | | | | | | | | | | | | | | | | |
|---|---|---|---|---|---|---|---|---|---|---|---|---|---|---|---|---|---|---|---|---|---|---|---|
| N | S | N | N | L | D | S | K | V | G | G | N | Y | N | Y | R | Y | R | L | F | R | K | S | N |
| P | P | P | P | N | A | P | B | N | N | N | P | P | P | P | B | P | B | N | N | B | B | P | P |
| P | P | P | P | H | O | P | E | H | S | S | P | H | P | H | E | H | E | H | H | E | E | P | P |

| | | | | | | | | | | | | | | | | | | | | | | | |
|---|---|---|---|---|---|---|---|---|---|---|---|---|---|---|---|---|---|---|---|---|---|---|---|
| L | K | P | F | E | R | D | I | S | T | E | I | Y | Q | A | G | S | K | P | C | N | G | V | E |
| N | B | N | N | A | B | A | N | P | P | A | N | P | P | N | N | P | B | N | P | P | N | N | A |
| H | E | S | H | O | E | O | H | P | P | O | H | H | P | H | S | P | E | S | S | P | S | H | O |

| | | | | | | | | | | | | | | | | | | | | | | | |
|---|---|---|---|---|---|---|---|---|---|---|---|---|---|---|---|---|---|---|---|---|---|---|---|
| G | F | N | C | Y | F | P | L | Q | S | Y | G | F | Q | P | T | N | G | V | G | Y | Q | P | Y |
| N | N | P | P | P | N | N | N | P | P | P | N | N | P | N | P | P | N | N | N | P | P | N | P |
| S | H | P | S | H | H | S | H | P | P | H | S | H | P | S | P | P | S | H | S | H | P | S | H |

##### Polarity features

| Polarity structure | Count | |
|---|---|---|
| | Reference | Mutation |
| ETC | 41 | 42 |

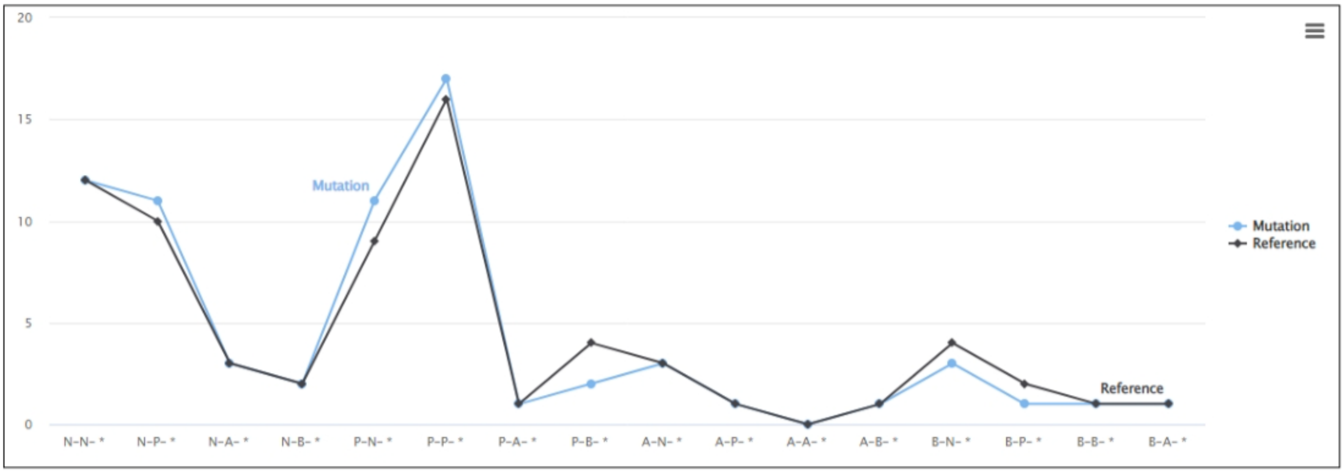

Amino acid polarity affects protein structure and stability. As a result, the amino acid polarity due to mutation of the amino acid sequence input by the user can be observed. We found repeated polarity patterns in the coronavirus and observed changes in the properties of amino acid sequence polarity due to mutation. Therefore, we provide visualization and table view of polarity pattern changes to the user.

① amino acid sequence
② 4 polarity characteristics
③ 5 amino acid properties
④ Mutated positions are indicated in red.

### Result report page – All viruses

#### Prediction Result

##### XBB

###### ① 3D Structure Prediction ⓘ

- The PAE is a value that estimates the difference between the relative locations of two residues of the model and the real model. The value has a negative correlation with the accuracy of the pairwise position of two residues. As the PAE value decreases, the accuracy increases and vice versa.
- The pLDDT is a value that estimates the reliability of the model. The value indicates the likelihood of folding of the protein structure at that location, with higher values indicating a greater likelihood of folding.

1 2 3 4 5 ⓘ Download all file

**Protein structure (Mutation)** Compare

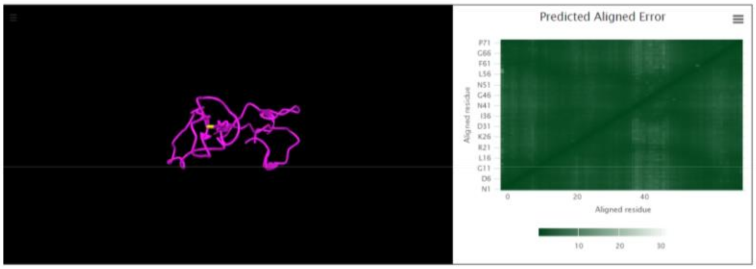

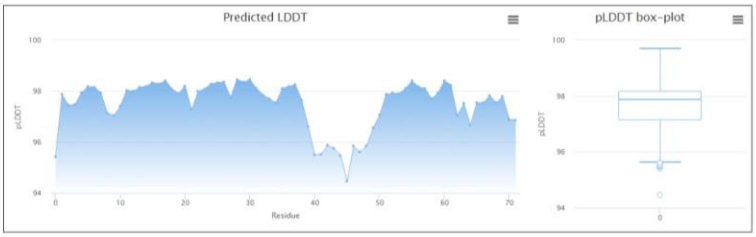

###### ② Virus amino acid info

- Visualizes and tabulates changes in repeated polarity structure sequence for wild-type sequences and mutated type sequences used in ①.

**Sequence without mutations (Reference)**

| | | | | | | | | | | | | | | | | | | | | | | | | | | | | | | | | | | | | | | | | | | | | | | | |
|---|---|---|---|---|---|---|---|---|---|---|---|---|---|---|---|---|---|---|---|---|---|---|---|---|---|---|---|---|---|---|---|---|---|---|---|---|---|---|---|---|---|---|---|---|---|---|---|
| N | S | N | K | L | D | S | K | P | S | G | N | Y | N | Y | L | Y | R | L | F | R | K | S | K | L | K | P | F | E | R | D | I | S | T | E | I | Y | Q | A | G | N | K | P | C | N | G | V | A |
| P | P | P | B | N | A | P | B | N | P | N | P | P | P | P | N | P | B | N | N | B | B | P | B | N | B | N | N | A | B | A | N | P | P | A | N | P | P | N | N | P | B | N | P | P | N | N | N |
| P | P | P | E | H | O | P | E | S | P | S | P | H | P | H | H | H | E | H | H | E | E | P | E | H | E | S | H | O | E | O | H | P | P | O | H | H | P | H | S | P | E | S | S | P | S | H | H |

| | | | | | | | | | | | | | | | | | | | | | | | |
|---|---|---|---|---|---|---|---|---|---|---|---|---|---|---|---|---|---|---|---|---|---|---|---|
| G | S | N | C | Y | S | P | L | Q | S | Y | G | F | R | P | T | Y | G | V | G | H | Q | P | Y |
| N | P | P | P | P | P | N | N | P | P | P | N | N | B | N | P | P | N | N | N | B | P | N | P |
| S | P | P | S | H | P | S | H | P | P | H | S | H | E | S | P | H | S | H | S | E | P | S | H |

**Sequence with mutations (Mutation)**

| | | | | | | | | | | | | | | | | | | | | | | | | | | | | | | | | | | | | | | | | | | | | | | | |
|---|---|---|---|---|---|---|---|---|---|---|---|---|---|---|---|---|---|---|---|---|---|---|---|---|---|---|---|---|---|---|---|---|---|---|---|---|---|---|---|---|---|---|---|---|---|---|---|
| N | S | N | K | L | D | S | K | P | S | G | N | Y | N | Y | L | Y | R | L | F | R | K | S | K | L | K | P | F | E | R | D | I | S | T | E | I | Y | Q | A | G | N | K | P | C | N | G | V | A |
| P | P | P | B | N | A | P | B | N | P | N | P | P | P | P | N | P | B | N | N | B | B | P | B | N | B | N | N | A | B | A | N | P | P | A | N | P | P | N | N | P | B | N | P | P | N | N | N |
| P | P | P | E | H | O | P | E | S | P | S | P | H | P | H | H | H | E | H | H | E | E | P | E | H | E | S | H | O | E | O | H | P | P | O | H | H | P | H | S | P | E | S | S | P | S | H | H |

| | | | | | | | | | | | | | | | | | | | | | | | |
|---|---|---|---|---|---|---|---|---|---|---|---|---|---|---|---|---|---|---|---|---|---|---|---|
| G | S | N | C | Y | S | P | L | Q | S | Y | G | F | R | P | T | Y | G | V | G | H | Q | P | Y |
| N | P | P | P | P | P | N | N | P | P | P | N | N | B | N | P | P | N | N | N | B | P | N | P |
| S | P | P | S | H | P | S | H | P | P | H | S | H | E | S | P | H | S | H | S | E | P | S | H |

| Polarity feature | | | Amino acid |
|---|---|---|---|
| Polarity features | Non-Polar | N | Ala (A), Val (V), Leu (L), Gly (G), Ile (I), Met (M), Trp (W), Phe (F), Pro (P) |
| | Polar | P | Ser (S), Cys (C), Asn (N) , Gln (Q), Thr (T), Tyr (Y) |
| | Acidic | A | Asp (D), Glu (E) |
| | Basic | B | Lys (K), Arg (R), His (H) |
| Five amino acid properties | Amino acids with electrically charged side chain_negative | 1 | Lys (K), Arg (R), His (H) |
| | Amino acids with electriccally charged side chain_positive | 2 | Asp (D), Glu (E) |
| | Amino acids with Polar uncharged side chain | 3 | Ser (S), Asn (N) , Gln (Q), Thr (T) |
| | Special cases | 4 | Cys (C), Gly (G), Pro (P) |
| | Amino acids with hydrophobic side chain | 5 | Ala (A), Val (V), Leu (L), Ile (I), Met (M), Trp (W), Phe (F) |

**Polarity features**

| Polarity structure | Count | |
|---|---|---|
| | Reference | Mutation |
| ETC | 45 | 45 |

### Result report page –Compare

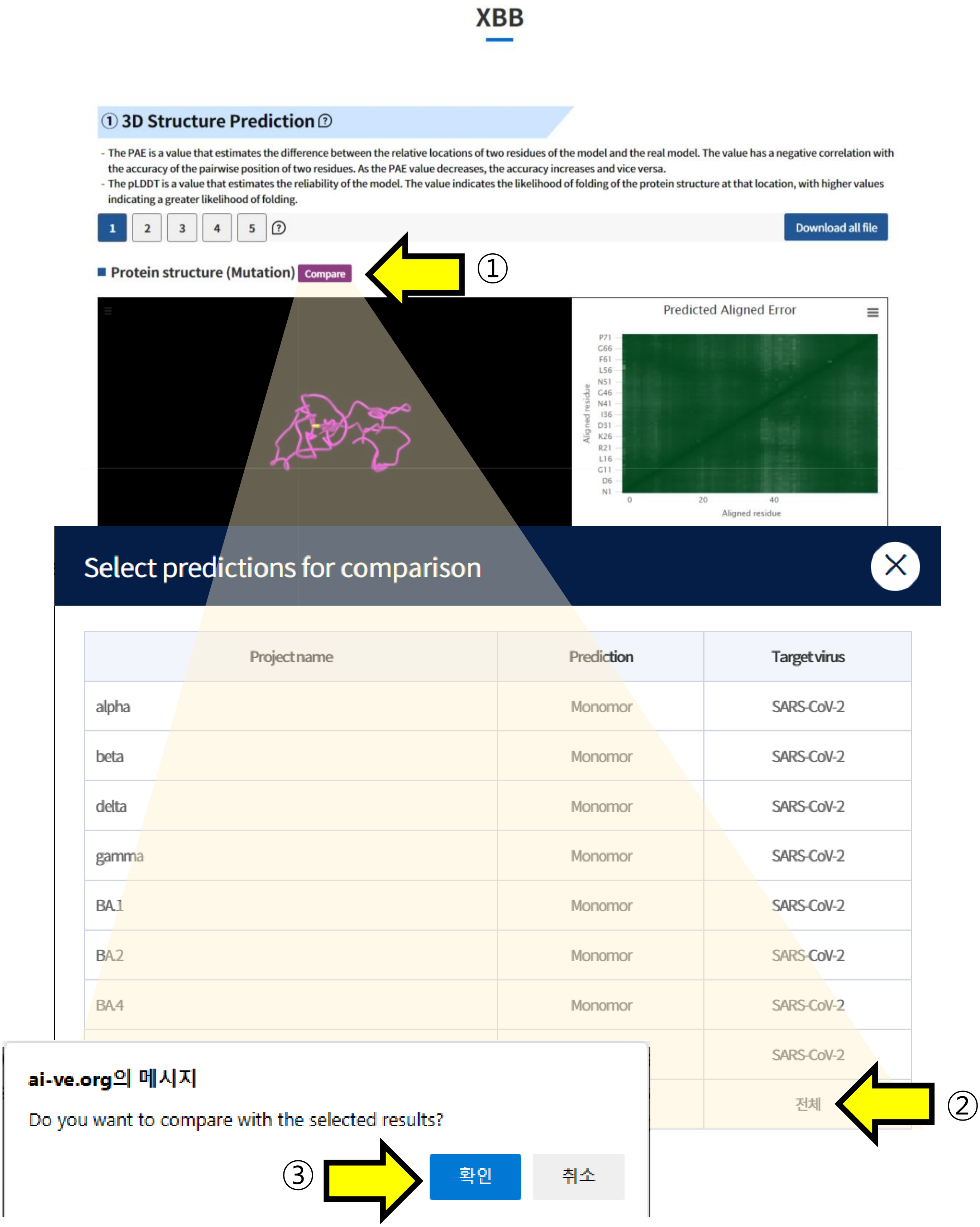

Structures predicted by “All viruses” can be compared with other structures using the compare feature.

① Click the "Compare" button to load the list of other tasks submitted by the user.

② Select the task you want to compare and click on it.

③ Click the "Confirm" button to navigate to the comparison page.

### Result report page –Compare

#### XBB compare to test

##### ① 3D Structure Prediction ⓘ

- The PAE is a value that estimates the difference between the relative locations of two residues of the model and the real model. The value has a negative correlation with the accuracy of the pairwise position of two residues. As the PAE value decreases, the accuracy increases and vice versa.
- The pLDDT is a value that estimates the reliability of the model. The value indicates the likelihood of folding of the protein structure at that location, with higher values indicating a greater likelihood of folding.

1 2 3 4 5 Download all file 1 2 3 4 5

■ Protein structure (Original)

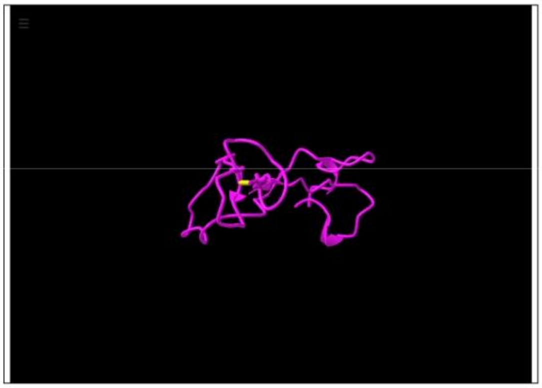

■ Protein structure (Compare target)

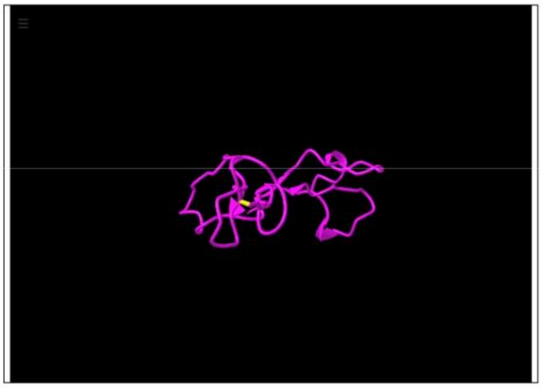

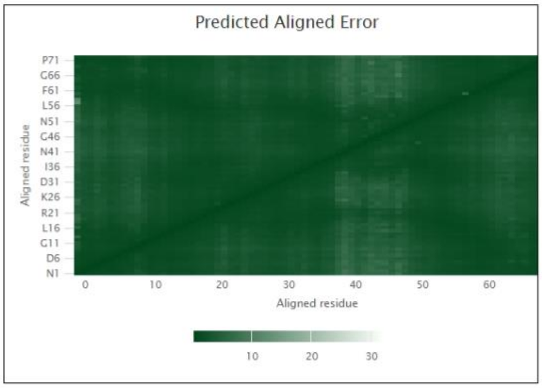

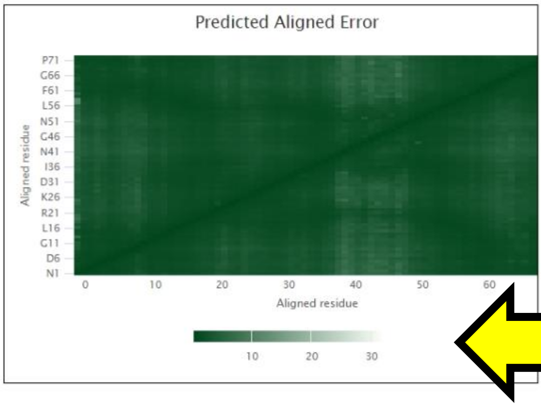

①

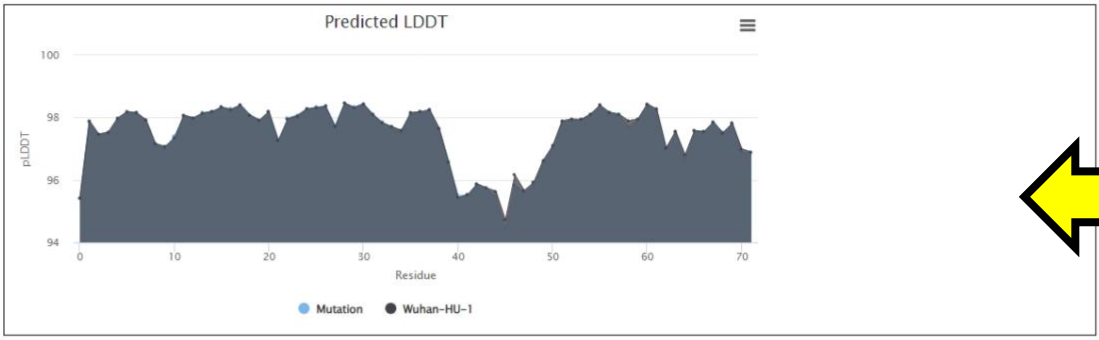

②

Through the "Compare" feature, you can compare two predicted structures in the list:
① You can visually inspect the PAE (Predicted Alignment Error) of the two structures using a plot.
② You can compare the pLDDT values of the two structures.

### Result report page –Compare

③

#### ② Virus amino acid info

- Visualizes and tabulates changes in repeated polarity structure sequence for wild-type sequences and mutated type sequences used in ①.

##### Sequence without mutations (Original)

| | | | | | | | | | | | | | | | | | | | | | | | | | | | | | | | | | | | | | | | | | | | | | | | |
|---|---|---|---|---|---|---|---|---|---|---|---|---|---|---|---|---|---|---|---|---|---|---|---|---|---|---|---|---|---|---|---|---|---|---|---|---|---|---|---|---|---|---|---|---|---|---|---|
| N | S | N | K | L | D | S | K | P | S | G | N | Y | N | Y | L | Y | R | L | F | R | K | S | K | L | K | P | F | E | R | D | I | S | T | E | I | Y | Q | A | G | N | K | P | C | N | G | V | A |
| P | P | P | B | N | A | P | B | N | P | N | P | P | P | P | N | P | B | N | N | B | B | P | B | N | B | N | N | A | B | A | N | P | P | A | N | P | P | N | N | P | B | N | P | P | N | N | N |
| P | P | P | E | H | O | P | E | S | P | S | P | H | P | H | H | H | E | H | H | E | E | P | E | H | E | S | H | O | E | O | H | P | P | O | H | H | P | H | S | P | E | S | S | P | S | H | H |

↓

| | | | | | | | | | | | | | | | | | | | | | | | |
|---|---|---|---|---|---|---|---|---|---|---|---|---|---|---|---|---|---|---|---|---|---|---|---|
| G | S | N | C | Y | S | P | L | R | S | Y | G | F | R | P | T | Y | G | V | G | H | Q | P | Y |
| N | P | P | P | P | P | N | N | B | P | P | N | N | B | N | P | P | N | N | N | B | P | N | P |
| S | P | P | S | H | P | S | H | E | P | H | S | H | E | S | P | H | S | H | S | E | P | S | H |

##### Sequence with mutations (Compare target)

| | | | | | | | | | | | | | | | | | | | | | | | | | | | | | | | | | | | | | | | | | | | | | | | |
|---|---|---|---|---|---|---|---|---|---|---|---|---|---|---|---|---|---|---|---|---|---|---|---|---|---|---|---|---|---|---|---|---|---|---|---|---|---|---|---|---|---|---|---|---|---|---|---|
| N | S | N | K | L | D | S | K | P | S | G | N | Y | N | Y | L | Y | R | L | F | R | K | S | K | L | K | P | F | E | R | D | I | S | T | E | I | Y | Q | A | G | N | K | P | C | N | G | V | A |
| P | P | P | B | N | A | P | B | N | P | N | P | P | P | P | N | P | B | N | N | B | B | P | B | N | B | N | N | A | B | A | N | P | P | A | N | P | P | N | N | P | B | N | P | P | N | N | N |
| P | P | P | E | H | O | P | E | S | P | S | P | H | P | H | H | H | E | H | H | E | E | P | E | H | E | S | H | O | E | O | H | P | P | O | H | H | P | H | S | P | E | S | S | P | S | H | H |

↓

| | | | | | | | | | | | | | | | | | | | | | | | |
|---|---|---|---|---|---|---|---|---|---|---|---|---|---|---|---|---|---|---|---|---|---|---|---|
| G | S | N | C | Y | S | P | L | Q | S | Y | G | F | R | P | T | Y | G | V | G | H | Q | P | Y |
| N | P | P | P | P | P | N | N | P | P | P | N | N | B | N | P | P | N | N | N | B | P | N | P |
| S | P | P | S | H | P | S | H | P | P | H | S | H | E | S | P | H | S | H | S | E | P | S | H |

| Polarity feature | | | Amino acid |
|---|---|---|---|
| Polarity features | Non-Polar | N | Ala (A), Val (V), Leu (L), Gly (G), Ile (I), Met (M), Trp (W), Phe (F), Pro (P) |
| | Polar | P | Ser (S), Cys (C), Asn (N) , Gln (Q), Thr (T), Tyr (Y) |
| | Acidic | A | Asp (D), Glu (E) |
| | Basic | B | Lys (K), Arg (R), His (H) |
| Five amino acid properties | Amino acids with electrically charged side chain_negative | 1 | Lys (K), Arg (R), His (H) |
| | Amino acids with electriccally charged side chain_positive | 2 | Asp (D), Glu (E) |
| | Amino acids with Polar uncharged side chain | 3 | Ser (S), Asn (N) , Gln (Q), Thr (T) |
| | Special cases | 4 | Cys (C), Gly (G), Pro (P) |
| | Amino acids with hydrophobic side chain | 5 | Ala (A), Val (V), Leu (L), Ile (I), Met (M), Trp (W), Phe (F) |

##### Polarity features

| Polarity structure | Count | |
|---|---|---|
| | Original | Compare target |
| ETC | 46 | 45 |

③ The sequence and polar structures of the two structures can be compared and analyzed.
